# Cholinergic control of GnRH neuron physiology and luteinizing hormone secretion: involvement of ACh/GABA co-transmission

**DOI:** 10.1101/2023.08.22.554236

**Authors:** Csaba Vastagh, Imre Farkas, Veronika Csillag, Masahiko Watanabe, Imre Kalló, Zsolt Liposits

**Author notes:** These authors contributed equally to this work (Csaba Vastagh, Imre Farkas). **Conflict of interest** The authors declare no competing financial interest.

## Abstract

Gonadotropin-releasing hormone (GnRH)-synthesizing neurons orchestrate reproduction centrally. Early studies have proposed the contribution of acetylcholine (ACh) to hypothalamic control of reproduction, although the causal mechanisms have not been clarified. Here, we report that *in vivo* pharmacogenetic activation of the cholinergic system increased the secretion of luteinizing hormone (LH) in orchidectomized mice. 3DISCO immunocytochemistry and electron microscopy revealed the innervation of GnRH neurons by cholinergic axons. Retrograde viral labeling initiated from GnRH-Cre neurons identified the medial septum and the diagonal band of Broca as exclusive sites of origin for cholinergic afferents of GnRH neurons. In acute brain slices, ACh and the ACh receptor (AChR) agonist carbachol evoked a biphasic effect on the firing rate in GnRH neurons, first increasing and then diminishing it. In the presence of tetrodotoxin, carbachol induced an inward current, followed by a decline in the frequency of mPSCs, indicating a direct influence on GnRH cells. RT-PCR and whole-cell patch-clamp studies revealed that GnRH neurons expressed both nicotinic (α4β2, α3β4, and α7) and muscarinic (M1-M5) ACh receptors. The nicotinic AChRs contributed to the nicotine-elicited inward current and the rise in firing rate. Muscarine via M1 and M3 receptors increased, while via M2 and M4 reduced the frequency of both miniature postsynaptic currents (mPSCs) and firing. Optogenetic activation of channelrhodopsin-2-tagged cholinergic axons modified GnRH neuronal activity and evoked co-transmission of ACh and GABA from a subpopulation of boutons. These findings confirm that the central cholinergic system immensely regulates GnRH neurons and activates the HPG-axis via ACh and ACh/GABA neurotransmissions.

**Significance statement:** Cholinergic drugs influence reproduction centrally, although the exact neuronal targets and regulatory mechanisms remain unsettled. We found that pharmacogenetic activation of the cholinergic system in vivo evoked an augmented LH release. The study also identified cholinergic cell groups in the mouse forebrain that innervate gonadotropin-releasing hormone (GnRH) neurons, the main hypothalamic regulators of reproduction. We also determined the subtypes of nicotinic and muscarinic receptors involved in neuronal information transmission and explored how their ligands affect the electrophysiological activity of GnRH neurons. A subset of cholinergic neurons was found to co-transmit GABA, which excites GnRH cells via GABA-A receptors. The findings suggest a novel, cholinergic regulation of the adult GnRH system in male mice that activates the pituitary-gonadal axis.

## Introduction

In vertebrates, the success of reproduction depends on the integrity of the hypothalamo-pituitary-gonadal (HPG) axis and the availability of the master molecule, gonadotropin-releasing hormone (GnRH) (Knobil, 1988). GnRH is synthesized in neurons of olfactory placode origin (Wray et al., 1989), which migrate through the basal forebrain and settle down at species-specific anatomical locations (Silverman et al., 1979; Merchenthaler et al., 1980; Liposits et al., 1984). Via their hypophysiotropic axon projections, the cells secrete GnRH in a pulsatile manner into the hypophysial portal circulation (Carmel et al., 1976) to control the proper gonadotroph hormone output of the anterior pituitary. The loosely organized GnRH cell population receives neuronal inputs from more than fifty different brain regions (Boehm et al., 2005), processes and integrates information on circulating gonadal hormones (Christian and Moenter, 2010; Farkas et al., 2018), and a wide range of peripheral metabolic signals (Farkas et al., 2013; Farkas et al., 2016; Csillag et al., 2019; Balint et al., 2020). Failure of cell migration, disturbances in sensing and processing peripheral hormone signals, or inability to respond properly to neurotransmitters and neuropeptides released from their synaptic afferents result in GnRH neuron and HPG axis dysfunctions or infertility (Pitteloud et al., 2006; Pitteloud et al., 2007). These pathological events interfere with the maintenance of species; therefore, the understanding of the diverse regulatory mechanisms of GnRH neurons at the molecular and cellular levels has been essential.

Most of the classic neurotransmitter systems, including glutamate (Iremonger et al., 2010) and GABA (DeFazio et al., 2002), have been shown to regulate reproduction centrally via synaptic channels in GnRH neurons. Acetylcholine (ACh) is a highly potent neurotransmitter used widely for communication in both the central and peripheral nervous systems. Its effects are executed via nicotinic (Albuquerque et al., 2009) and muscarinic ACh receptor (Caulfield and Birdsall, 1998) subtypes that are differentially expressed in diverse neuron phenotypes. Cholinergic signaling has been proposed to control reproduction centrally. Early *in vivo* studies have shown that nicotine delays the surge release of luteinizing hormone (LH) (Blake et al., 1972), inhibits the progesterone-advanced LH surge in proestrus (Kanematsu and Sawyer, 1973), blocks the LH secretion induced by opioid receptor antagonists (Hodson et al., 1997), and decreases the pulsatile secretion of LH (Sano et al., 1999). Under *in vitro* conditions, the natural ligand ACh stimulated the release of GnRH from hypothalamic fragments and the secretion of LH (Fiorindo and Martini, 1975). The involvement of muscarinic mechanisms has also been shown since selective muscarine receptor antagonists stimulated the release of GnRH from the median eminence of the rat (Koren et al., 1992). The use of immortalized GnRH neurons (GT1-7 cells) allowed the characterization of some molecular mechanisms involved in the cholinergic modulation of GnRH neurons. These cells express functional, high-affinity α4β2 and α7 subtypes of nicotinic ACh receptors (nAChR), and the activation of the α-bungarotoxin-sensitive α7 subclass by nicotine exerts an inhibitory effect on GnRH release (Messi et al., 2018). A biphasic action of ACh on GnRH release was seen in perifused hypothalamic cells and GT1-7 neurons, whereas nicotine alone increased and muscarine decreased the release of the neurohormone (Krsmanovic et al., 1998). While nicotinic and muscarinic M1 receptors exert stimulatory effects, activation of muscarinic M2 receptors inhibits GnRH release (Krsmanovic et al., 1998). Transcriptome profiling of rodent GnRH neurons and GT1-7 cells has confirmed the expression of certain nicotinic and muscarinic receptors (Todman et al., 2005; Varju et al., 2009; Arai et al., 2017), supporting the view of a direct action of cholinergic drugs upon GnRH neurons. It has been shown recently that murine GnRH neurons grown in nasal explants receive regulatory acetylcholine neurotransmission (Shostak et al., 2023). Although these findings unequivocally imply that the central cholinergic system is capable of modulating GnRH neurons, the molecular, cellular, and physiological features of this regulatory channel are largely unidentified in adult mice.

Therefore, the present study was undertaken in adult male mice to (1) measure the impact of *in vivo* chemogenetic activation of cholinergic neurons on luteinizing hormone secretion; (2) explore the putative networking of the central cholinergic system with hypophysiotropic GnRH neurons; (3) elucidate the source(s) of ACh involved in neurotransmission; (4) reveal the type(s) of cholinergic receptors expressed in GnRH neurons, and (5) characterize the effects of ACh receptor ligands on the physiology of GnRH cells.

## Materials and Methods

### Ethics

All experiments were performed in accordance with the Institutional Ethical Codex and the Hungarian Act of Animal Care and Experimentation Guidelines (40/2013, II.14), which are in concert with the European Communities Council Directive of September 22, 2010 (2010/63/EU). The Animal Care and Experimentation Committee of the Institute of Experimental Medicine and the National Scientific Ethical Committee on Animal Experimentation have also approved the experiments under the project numbers PE/EA/796-5/2021 and PE/EA/00646-2/2022.

### Animals

Adult male wild-type, and transgenic mice on the C57Bl/6 J or Bl6Fx background were used from local colonies bred at the Medical Gene Technology Unit of the Institute of Experimental Medicine (IEM). All animals were housed in a light (12:12 h light–dark cycle, lights on at 06:00 h) and temperature (22 ± 2 °C) controlled environment, with free access to standard food and tap water. All surgical interventions and perfusions were carried out under deep anesthesia by intraperitoneal injections of a cocktail of ketamine (25 mg/kg body weight) and xylazine (5 mg/kg body weight).

### Transgenic mouse lines and characterizations

#### The GnRH-*GFP* mouse line

The GnRH-GFP transgenic mouse line was generated to facilitate the study of hypothalamic GnRH neurons in which the green fluorescent protein (GFP) is genetically targeted to these cells (Suter et al., 2000b). The expression of GFP was detected in 84–94% of GnRH immune-positive neurons. This mouse line was used in studies aimed at electrophysiological characterization (Suter et al., 2000a; Christian et al., 2005; Pielecka-Fortuna et al., 2008; Farkas et al., 2010; Farkas et al., 2016) and cell type-specific genomic analysis of the GnRH neurons (Todman et al., 2005; Vastagh et al., 2015; Vastagh et al., 2016).

#### The Chat-*Cre* mouse line

Chat-IRES-Cre knock-in mice that express Cre recombinase in cholinergic neurons were bought from the Jackson Laboratory (JAX stock #006410). The line was re-derived at the Medical Technology Unit of the IEM, Hungary (Van Keuren and Saunders, 2004). Stock #006410 animals were bred to the ROSA26::FLPe strain to facilitate the removal of the *frt*-neo cassette between the Cre and SV40 poly(A) to exclude ectopic Cre expression. The resulting animals with the Chat-Cre-Δneo knock-in allele were bred together, while the ROSA26::FLPe was removed, to establish the chat/cre_Δneo//Gt(ROSA)26Sor/CAG/LSL mouse line.

#### The *Chat*-Cre-ChR2 mouse line

Chat-Cre-Δneo male mice were crossed with channelrhodopsin-2(ChR2) reporter mice [Ai32(RCL-ChR2(H134R)/EYFP), JAX stock #012569, for the purpose of optogenetics and morphological studies (Madisen et al., 2012). Using an eyfp reference gene assay mix, the copy number of the Cre and enhanced yellow fluorescent protein (eYFP) transgenes in the offspring was measured. Animals homozygous for Cre and eYFP were selected for morphological experiments as well as for crossbreeding with the GnRH-GFP mouse line. The cholinergic neuron-specific expression of ChR2 was confirmed by double immunofluorescence staining for the choline acetyltransferase (Chat) enzyme and eYFP.

#### The Chat-Cre-ChR2-GnRH-*GFP* mouse line

The triple-transgenic, Chat-Cre-ChR2-GnRH-GFP mouse line was established for the purpose of the optogenetic study using *in vitro* acute brain slice electrophysiology. In this experimental design, blue LED (470 nm) light stimulation of the ChR2-expressing axons of cholinergic neurons distributed in the medial preoptic area and whole-cell patch recording from GFP-expressing GnRH neurons were performed simultaneously. To carry out this goal, homozygous male Chat-Cre-ChR2 mice were crossed with female GnRH-GFP mice. The presence of the GnRH-GFP transgene in the offspring of the first generation was confirmed using real-time qPCR; the relative copy number of the gfp transgene (3 vs. 2) was distinguishable using this method. The immunocytochemical phenotyping confirmed that GnRH neurons receive innervation from channelrhodopsin-2-expressing cholinergic neurons.

#### The Chat-Gq (hM3Dq) *DREADD* mouse line

Chat/Cre_Δneo mice were crossed with B6N;129-Tg(CAG-CHRM3*,-mCitrine)1Ute/J mice (The Jackson Laboratories, JAX stock #026220) to express designer receptors (DREADD) signaling via Gq (hM3Dq) (Armbruster et al., 2007) in cholinergic neurons. This mouse line was used for pharmacogenetics. Its immunocytochemical characterization revealed mCitrine expression in Chat-immunoreactive neurons of the brain. The established new mouse line was used for *in vivo* chemogenetic activation of the cholinergic system, including those subsets of cholinergic neurons that modulate the hypothalamic control mechanisms of reproduction. The immunocytochemical phenotyping of the transgenic mouse strain confirmed the cholinergic neuron-specific expression of the excitatory hM3Dq DREADD receptor.

#### The GnRH-Cre *mouse* line

Transgenic mice expressing Cre recombinase in GnRH neurons (GnRH-Cre) were used to determine brain regions that host cholinergic cells innervating GnRH neurons. The generation and characterization of this mouse line were published elsewhere (Yoon et al., 2005). Briefly, a 212 kb bacterial artificial chromosome (BAC) holding the entire *GnRH* coding sequence was modified: a *Cre recombinase-poly(A)* cassette was inserted into the ATG of the first exon of the *GnRH* gene, resulting in a high (over 96 percent) co-localization rate of the anti-Cre and anti-GnRH immunoreactivities (Yoon et al., 2005).

#### Pharmacogenetics

Transgenic male (n = 13) mice expressing mCitrine and activating (hM3Dq) designer receptors exclusively activated by designer drugs (DREADDs) (chat/cre_Δneo//GT(ROSA)26Sor_CAG/LSL_CHRM3_mCitrine (TgTm): Bl6Fx) in cholinergic neurons were used. Mice were bilaterally orchidectomized (ORX) under ketamine-xylazine anesthesia. After a week of recovery, all mice were handled and trained for tail-tip bleeding every day (McCosh et al., 2018), for at least 28 days. A week before blood collection, animals were also trained every day to lick off a small bolus of low sugar containing peanut cream (∼20mg) mixed with 1µl DMSO after stroking the tail (Mamgain et al., 2021).

After clipping the tip of the tail (1-2 mm), mice were placed on a flat surface every 3 minutes and allowed to roam freely while 3 µl of blood was collected from the tail wound. The first seven to ten blood samples were discarded to eliminate samples deriving from the after-cut period potentially affected by stress. The subsequent blood samples were immediately mixed with 57 µl of 0.1 M PBS (Invitrogen) containing 0.05% Tween 20 and 0.2% BSA (Jackson ImmunoResearch) and preserved in prediluted form in 60 µl aliquots on ice during blood collection, then stored at −20°C until LH assay. Samples for LH measurements were collected for 180 minutes, which generated 60 aliquots per mouse. After collecting the 29^th^ blood sample, animals either received vehicle (1 µl DMSO; n = 6) or the DREADD ligand clozapine n-oxide (CNO, Tocris; 0.1 mg/µl DMSO; 5 mg/kg body weight; n = 7) in a single bolus of peanut cream.

#### Ultrasensitive luteinizing hormone (LH) measurement

The secretion pattern of luteinizing hormone (LH) secretion was monitored and measured by an extremely sensitive ELISA technique (Steyn et al., 2013) developed a decade ago. In the experiment, the intra- and inter-assay coefficients of variation were 5.54 ± 2.15% and 7.62 ± 1.33%, respectively. Overall basal levels of LH secretion were determined by combining the 10 lowest LH measurements from each mouse (Czieselsky et al., 2016). The mean levels of LH release were the average of all values collected over the sampling period. Basal and mean LH levels before and after administration of CNO or vehicle were determined and analyzed between treatment periods using paired t test.

### Immunocytochemistry

#### Fluorescent immunocytochemistry

Adult, intact (n = 15) and transgenic mice (n = 20) were perfused transcardially with 4% paraformaldehyde (pH = 7.6 in PBS) for 10 min at a flow rate of 4 ml/min. Brains were removed rapidly from the skulls and post-fixed in the same fixative overnight at 4 C. On the next day, brains were sectioned in the coronal plane at 25 μm thickness using a vibratome (Leica Inc., Wetzlar, Germany). Sections containing the medial septum (MS), diagonal band of Broca (DBB), and medial preoptic area (mPOA) were pre-treated with 0.5% H_2_O_2_ and permeabilized with 0.5% Triton X-100 for 20 min. The following primary antisera were used: anti-GnRH [raised in guinea pig #1018, diluted in 1:10.000 (gift from E. Hrabovszky)], anti-GFP (raised in either rabbit [Millipore #AB10145, 1:2.000 dilution]), and anti-VAChT (raised in rabbit, 1:5000 dilution) anti-vesicular GABA transporter (VGAT, raised in goat, 1:2500 dilution), and anti-Chat (Millipore #AB144P, raised in goat, 1:500 dilution). Secondary antisera (Alexa 488 conjugated anti-rabbit IgG; Cy3 conjugated anti-goat IgG; Cy5 conjugated anti-guineapig [Jackson ImmunoResearch Europe Ltd, UK]) were all diluted at 1:1000 in serum diluent (2% normal horse serum in PBS).

#### 3DISCO technique

Adult male GnRH-GFP transgenic mice (n = 6) were perfused, and the brains were post-fixed as described in the previous section. Brains placed in a mouse brain mold were sliced at 1 mm thickness in the sagittal plane. Thereafter, double fluorescent immunocytochemistry combined with the 3DISCO technique (Ertürk and Şengül, 2012) was applied (Vastagh et al., 2021). Briefly, the slices were rinsed several times in PBS, then permeabilized with 0.5% Triton-X100 in PBS containing 0.1% gelatin and 0.05% merthiolate (PBS-GT) for 3 days at room temperature (RT) under constant agitation. For fluorescent immunocytochemical double labeling, brain slices were incubated with the anti-GFP (goat; #AB5450 Abcam, Cambridge, UK) and anti-VAChT (rabbit; generated by M. Watanabe, Hokkaido University, Japan) primary antibodies diluted in 1:5000 in PBS-GT for 7 days at RT. Several washes (6 x 60 mins) in PBS-GT were followed by the incubation with secondary antibodies (1:500; Cy3 anti-goat IgG and Cy5 anti-rabbit IgG; Jackson ImmunoResearch Europe Ltd., Cambridge House, UK) for 7 days at RT.

Slices were then rinsed in PBS-GT for 6 x 60 mins and then in PBS for 30 minutes. For dehydration and lipid removal steps, a series of increasing tetrahydrofuran (THF, Sigma-Aldrich) concentrations were used (50% THF overnight at 4°C, 80 %, and 3 x 100% THF for 30 min at RT). For optical clearing, slices were immersed in dibenzyl ether (DBE, Sigma-Aldrich) for at least 15 minutes, then kept either in DBE for storage or covered using a glass slide with custom-made plastic supporting frames adjusted for the actual specimen thickness. The specimen was cover-slipped with DBE for the later confocal microscopic analysis.

#### Quantitative 3D image analysis of the interacting cholinergic and GnRH neuronal systems

For quantitative evaluation of the interacting cholinergic and GnRH neuronal systems, stacks of images were acquired with a Nikon C2 laser scanning confocal system (Nikon Instruments Europe BV, Amsterdam, Netherlands). To maximize the emitted signal intensity, sequential scanning mode was set in the software (NIS Elements AR, Nikon). For Cy3-coupled labeling, an argon laser (561 nm) and a 595/40 nm emission filter (Nikon) were used. At the same time, to detect Cy5 emission, a He-Ne laser (637 nm) and a DAPI/CY5 (custom #239415 Semrock, Lake Forest, USA) dual emission filters were applied. The sampling of interacting neuronal profiles was carried out in 2 slices per mouse brain and 4 adjacent frames of each sagittal brain slice. A total of 48 imaging areas were studied in 6 brains that contained the residence of GnRH neurons (see **Fig. 2**).

For quantitative analysis, 20x objective magnification (Plan-Apo VC DIC N2, NA: 0.75 WD: 1mm) was used at 1024 x 1024-pixel *x-y* image resolution. During the scanning process, the pinhole diameter was set to 1 AU (Airy unit) in the acquisition software. The optical thickness was 2 µm. The optical sectioning was adjusted to match pinhole size (z-step interval: 1.2 µm).

All GFP immunoreactive (IR) perikarya were counted in the z-stacks of scanned areas. In the orthogonal view of the z-stacks, GFP-IR neurons that received VAChT-IR axonal appositions were also counted. The ratio of GFP-IR neurons targeted by VAChT axons was calculated. The ratio of all GnRH-IR dendrites to those contacted by VAChT-IR axons was estimated similarly. Student’s t-test was used to test the significance of the differences in the morphological data and was considered significant at p < 0.05.

#### Immunoelectron microscopy

GnRH-GFP transgenic mice (n = 10) were anesthetized and perfused transcardially with 10 ml of 0.01 M phosphate-buffered saline (PBS), pH 7.4, followed sequentially by 30 ml of fixative containing 4% acrolein and 2% paraformaldehyde (PFA) in PBS, pH 7.4, and then by 40 ml of 4% PFA in PBS, pH 7.4. The brains were removed and stored in 4% PFA in 0.1M PB overnight at 4 °C. Serial 25-µm-thick, coronal sections were cut on a Leica VT 1000S vibratome (Leica Microsystems, Vienna, Austria) through the medial septum-diagonal band of Broca and the medial preoptic area. The sections were treated with 1% sodium borohydride in 0.1M PB for 30 minutes, followed by 0.5% H_2_O_2_ in PBS for 15 minutes, cryoprotected in 15% sucrose in PBS for 15 minutes at room temperature, and in 30% sucrose in PBS overnight at 4°C. The sections were incubated with 2% normal horse serum for 20 minutes to block the non-specific binding of antibodies. The tissue penetration of antibodies was facilitated with Photo-Flo (Kodak Photo-Flo 200 Solution) added to the used antibodies at a concentration of 0.3%.

The pre-treated sections were placed in a mixture of guinea pig anti-GFP serum (1:2500) and anti-VAChT serum (1:10.000) for 4 days at 4 °C. After rinsing in PBS and 0.1% cold water fish gelatin and 1% bovine serum albumin (BSA) in PBS, the sections were incubated in donkey anti-sheep IgG conjugated with 0.8 nm colloidal gold (Electron Microscopy Sciences, Fort Washington, PA) diluted at 1:100 and biotinylated anti-rabbit IgG diluted at 1:500 in PBS containing 0.1% cold water fish gelatin and 1% BSA overnight at 4°C. After washing in PBS, the sections were fixed in 1.25% glutaraldehyde in 0.1M PB for 10 min at room temperature. After further rinsing in PBS, the sections were washed in Aurion ECS buffer (Aurion, Wageningen, Netherlands) (1:10, diluted in distilled water. The sections were further rinsed in 0.2 M sodium citrate pH 7.5, and then the gold particles were silver-intensified with the Aurion R-Gent SE-LM Kit. The sections were placed in 0.05% gold-chloride for 2×5 min at room temperature, washed in 0.2 M sodium citrate, pH 7.5, and in 3% sodium-thiosulphate solution for 10 min each at room temperature. Then the sections were treated in avidin/biotin-peroxidase complex (Vectastain Elite ABC Elite 1:500, Vector Laboratories Inc, Burlingame, US) and the GnRH-immunoreactivity was developed in 0.05% DAB /0.005% H_2_O_2_ in 0.05 M Tris buffer, pH 7.6.

The sections were processed for electron microscopy, first osmicated for 30 minutes (OsO_4,_ 1%) at room temperature, and then treated with 2% uranyl acetate in 70% ethanol for 30 min. Following dehydration in an ascending series of ethanol and propylene oxide (Sigma), the sections were flat embedded in Araldite 6005 epoxy resin (EMS) on liquid release agent-coated slides and polymerized at 56 °C for 2 days. After polymerization, 60–70 nm-thick ultrathin sections were cut with a Leica UCT ultramicrotome (Leica Microsystems, Wetzlar, Germany). The ultrathin sections were mounted onto Formvar-coated, single-slot grids and examined with a Hitachi H-7100 transmission electron microscope (Hitachi Ltd., Tokyo, Japan).

### Cre-dependent retrograde rabies virus tract tracing

#### Stereotaxic surgery

A cocktail of helper viruses (pAAV-EF1a-FLEX-TVA-mCherry /pAAV-CAG-FLEX-oG-WPRE-SV40pA in a ratio of 1:1, Salk Institute, La Jolla) was delivered by a single injection into the MS/DBB region (Bregma antero-posterior +0.85 mm, dorso-ventral −4.5 mm, and medio-lateral: 0 mm) and bilateral injections into the MPA (Bregma antero-posterior +0.49 mm, dorso-ventral −5 mm, and medio-lateral: ±0.25 mm) of male GnRH-Cre mice (n = 15). Three weeks later, bilateral injections were made with the rabies ΔG-EnvA-eGFP virus (BRVenvA-2C, Charité-Universitätsmedizin Berlin, Viral Core Facility) into the helper virus-infiltrated territory (Bregma antero-posterior +0.6 mm, dorso-ventral −5 mm, and medio-lateral: ±0.25 mm). Due to the expression of the transgenes, helper virus-infected cells became red (in a Cre-dependent manner), while rabies virus-infected cells became green (at the injection site, in a random manner). The starter cells expressing both transgenes became yellow. The helper virus-infected cells contained an upgraded version of the rabies glycoprotein (oG) that has increased the trans-synaptic labeling potential (Kim et al., 2016). After 8 - 14 days of survival, mice were perfused with 4% PFA, brains were removed, and 30 µm-thick sections were cut on a freezing microtome (Reichert).

#### Immunohistochemical identification of rabies-infected cholinergic neurons

The fluorescent marker proteins mCherry and GFP expressed in response to the viral infections and the cholinergic marker enzyme Chat were detected by a triple-label immunofluorescent technique using rabbit anti-DS Red IgG (Clontech, #632496), chicken anti-GFP (Invitrogen, #A10262), and goat anti-Chat (Millipore, #AB144P) primary antibodies, and CY3, FITC, and CY5 conjugated anti-rabbit (Jackson Lab, #715-165-152), anti-chicken (Jackson Lab, #703-095-155) and anti-goat (Jackson Lab, #705-165-147) secondary antibodies, respectively. Sections were mounted, cover slipped by Prolong Gold Antifade, and scanned by a slide scanner (Pannoramic Scan; 3D-HISTECH, Budapest, Hungary) using 10x objective magnification. Selected sections underwent confocal analyses using a Nikon A1 confocal microscope at 20x and 60x magnification.

#### Immunohistochemical controls

The specificity of the primary antisera used has been reported previously. Accordingly, increasing dilutions of the primary antibodies resulted in a gradual decrease and eventual disappearance of the immunostaining; omission of the primary antibodies or their pre-absorption with corresponding peptide antigens resulted in a complete loss of the immunostaining. The secondary antibodies used here were designed for multiple labeling and were preabsorbed by the manufacturer with IGs from several species, including the one in which the other primary antibody had been raised.

### Real-time polymerase chain reaction (RT-PCR)

#### Cytoplasm sampling

GnRH cytoplasmic samples were harvested from acute forebrain slices of adult male mice containing GnRH-GFP neurons using a patch pipette. A total of 7 samples were processed, with 10 cytoplasmic fractions per sample. The tips of the glass pipettes were broken into PCR tubes, and then cDNA was synthesized from the sample RNA using SuperScript IV VILO Master Mix (#11766050, Thermo Fisher Scientific, Waltham, US) according to the manufacturer’s instructions.

#### PCR

Preamplification ran for 14 cycles (TaqMan PreAmp Master Mix Kit, Thermo Fisher Scientific, Waltham, US). Gene expression analyses were performed on a ViiA 7 real-time PCR system (Thermo Fisher Scientific) using TaqMan assays (Thermo Fisher Scientific) according to the manufacturer’s instructions for the following target genes: *Gapdh, Hprt, GnRH1, Chrm1, Chrm2, Chrm3, Chrm4, Chrm5, Chrna2, Chrna3, Chrna4, Chrna5, Chrna6, Chrna7, Chrnb1, Chrnb2, Chrnb3, Chrnb4, Chrnd, Chrne* and *Chrng*. The real-time PCR experiments were programmed to run for 40 cycles. Expression of the *Gnrh* gene was detectable in all 7 cell pools. The mean expression of *Gapdh* and *Hprt* genes in each of the cytoplasmic sample pools (CPs) was used as a reference (Ct _ref_). Gene expression level (G) of the target genes was calculated as follows: G = 2 ^-ΔCt^, where ΔCt is the difference in threshold cycle between the target and reference genes (ΔCt = Ct _gene_-Ct _ref_) (Riedel et al., 2014).

### Electrophysiology

#### Slice electrophysiology

Brain slice preparation was carried out as described earlier (Farkas et al., 2010). Briefly, after decapitation, the heads were immersed in ice-cold, low-Na cutting solution and continuously bubbled with carbogen, a mixture of 95% O_2_ and 5% CO_2_, and the brains were removed rapidly from the skull. The cutting solution contained the following (in mM): saccharose 205, KCl 2.5, NaHCO_3_ 26, MgCl_2_ 5, NaH_2_PO_4_ 1.25, CaCl_2_ 1, and glucose 10. Hypothalamic blocks were dissected, and 220-μm-thick coronal slices were prepared from the mPOA with a VT-1000S vibratome (Leica Microsystems, Wetzlar, Germany) in the ice-cold, low-Na, oxygenated cutting solution. The slices containing preoptic area (POA) were transferred into artificial cerebrospinal fluid (aCSF) (in mM): NaCl 130, KCl 3.5, NaHCO_3_ 26, MgSO_4_ 1.2, NaH_2_PO_4_ 1.25, CaCl_2_ 2.5, and glucose 10, bubbled with carbogen, and left for 1 h to equilibrate. Equilibration started at 33^◦^C, and it was allowed to cool down to room temperature.

Recordings were carried out in carbogenated aCSF at 33^◦^C. Axopatch-200B patch-clamp amplifier, Digidata-1322A data acquisition system, and pCLAMP 10.4 software (Molecular Devices Co., Silicon Valley, CA, USA) were used for recording. Neurons were visualized with a BX51WI IR-DIC microscope (Olympus Co., Tokyo, Japan). The patch electrodes (OD = 1.5mm, thin wall; WPI, Worcester, MA, USA) were pulled with a Flaming-Brown P-97 puller (Sutter Instrument Co., Novato, CA, USA).

GnRH-GFP neurons were identified by brief illumination at 470 nm using an epifluorescent filter set based on their green fluorescence, typical fusiform shape, and characteristic topography (Suter et al., 2000b).

#### Whole-cell patch clamp

Whole-cell patch-clamp measurements started with a control recording (1 - 5 min), then the selected receptor ligand was pipetted into the aCSF-filled measurement chamber containing the brain slice in a single bolus, and the recording continued for a further 7 - 10 min. Pre-treatment with the extracellularly applied antagonist started 15 minutes before adding the ligand, and the antagonist was continuously present in the aCSF during the electrophysiological recording. Each neuron served as its own control when drug effects were evaluated. The membrane current and the mPSCs in GnRH neurons were measured as described earlier (Farkas et al., 2010). Briefly, the neurons were voltage clamped at −70 mV of holding potential. The intracellular pipette solution contained the following (in mM): HEPES 10, KCl 140, EGTA 5, CaCl_2_ 0.1, Mg-ATP 4, and Na-GTP 0.4 (pH = 7.3 with NaOH). The resistance of the patch electrodes was 2–3 MΩ. Only cells with aa low holding current (10 pA) and a stable baseline were used. Input resistance (R_in_), series resistance (R_s_), and membrane capacitance (C_m_) were also measured before and after each treatment by using 5 mV hyperpolarizing pulses. To ensure consistent recording qualities, only cells with R_s_ <20 MΩ, R_in_ >500 MΩ, and C_m_ >10 pF were accepted. Spike-mediated transmitter release was blocked in all mPSC experiments by adding the voltage-sensitive Na-channel inhibitor tetrodotoxin (TTX; 660 nM, Tocris) to the aCSF 10 min before mPSCs were recorded. Time distribution graphs of frequencies were generated using 30 s time bins, shifted by 5 s steps, to show the time courses of the effect of substances.

Action potentials were recorded in whole-cell current clamp mode at 0 pA. After a control period (1-3 min), the agonists were applied, and the recording continued for 8 - 10 min. The antagonist pretreatments started 10 minutes before starting the recording. Each neuron served as its own control when drug effects were evaluated.

The intracellularly applied tetrahydrolipstatin (THL, Tocris, 10 μM) was added to the intracellular (pipette) solution. After achieving the whole-cell configuration, measurements started after 15 minutes of equilibration to reach a stable intracellular milieu.

Statistical significance was analyzed by Student’s t-test and one-way ANOVA, followed by Tukey’s post-hoc test on the percentage and on the current amplitude data, and considered significant at p < 0.05.

#### Optogenetics

The applied technique has recently been published (Vastagh et al., 2021) with a slight modification. A LED light source (CoolLED pE-100, Andover, UK, 470 nm) was fitted to the microscope, illuminating the sample via the objective lens. Whole-cell patch clamp measurements were carried out to record mPSCs in voltage-clamp or action potentials (APs) in current-clamp mode in GnRH neurons in acute brain slices of triple-transgenic Chat-Cre-ChR2-GnRH-GFP adult male mice. The neurons were voltage clamped at a pipette holding potential −70 mV (for PSCs) and current clamped at 0 pA (for APs). The duration of a LED pulse was 5 ms.

For evoked mPSCs (emPSC), LED pulses were applied at 0.2 Hz or 5 Hz (60 runs total; each run was 5 s long), and then records of the emPSC responses of the 60 runs were averaged (9 neurons from 5 mice). The AChR receptor antagonists mecamylamine and atropine, or the GABA_A_-R blocker picrotoxin were added to the aCSF, and 10 minutes later, the measurement was repeated.

Measurement of the firing rate or mPSC frequency changes was carried out using LED pulses. The measurements started with a recording with no LED (for control purposes, 30 s for APs, 2 min for mPSCs) and with a subsequent LED train of 5 Hz illumination period (60 s for APs, 2 min for mPSCs). Analysis of the frequency changes was carried out on the percentage data calculated by dividing the frequency of the second period with that of the first one (control period) on 10 recorded cells from 5 animals. The ACh receptor antagonists mecamylamine and atropine were added to the aCSF, and 10 minutes later, the measurement was carried out in the presence of the antagonists.

#### Data analysis

Recordings were stored and analyzed offline. Event detection was performed using the Clampfit module of the PClamp 10.4 software (Molecular Devices Co., Silicon Valley, CA, USA). Firing rates and mPSC frequencies were calculated as the number of APs or mPSCs divided by the length of the corresponding time. The mean values of the control and treated periods of the recording are calculated from these frequency values. All the experiments were self-controlled in each neuron: percentage changes in the firing rate or frequency of the mPSCs were calculated by dividing the frequency value in the treated period with that of the control period. Group data were expressed as mean ± standard error of mean (SEM). Two-tailed Student’s t-test, or ANOVA was applied for comparison of groups, and the differences were considered significant at p < 0.05. The detailed data for statistical analyses are in **Tables 2-4**.

**Table 1.**
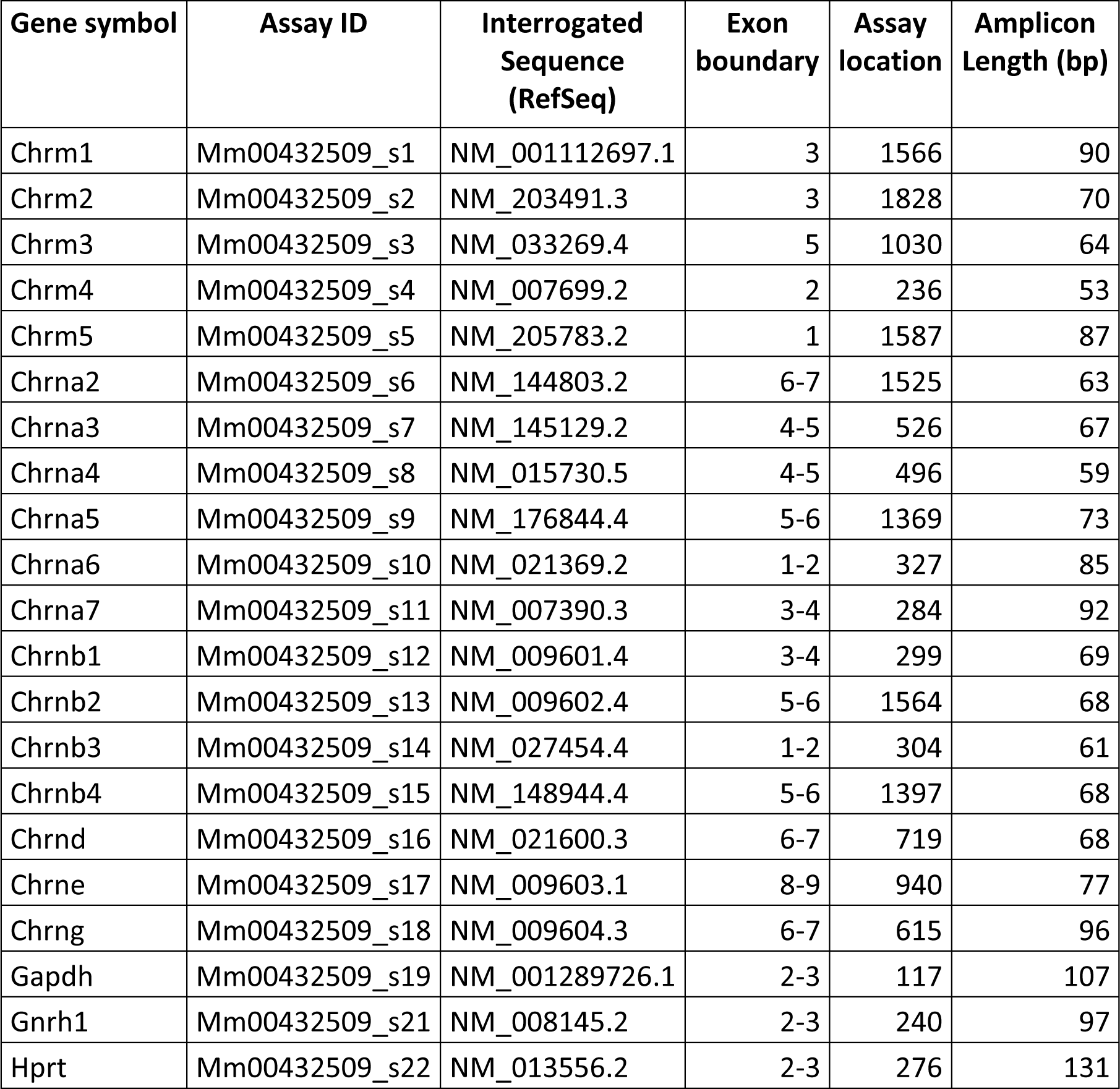
Primer probes used for the quantitative RT-PCR study.

**Table 2.**
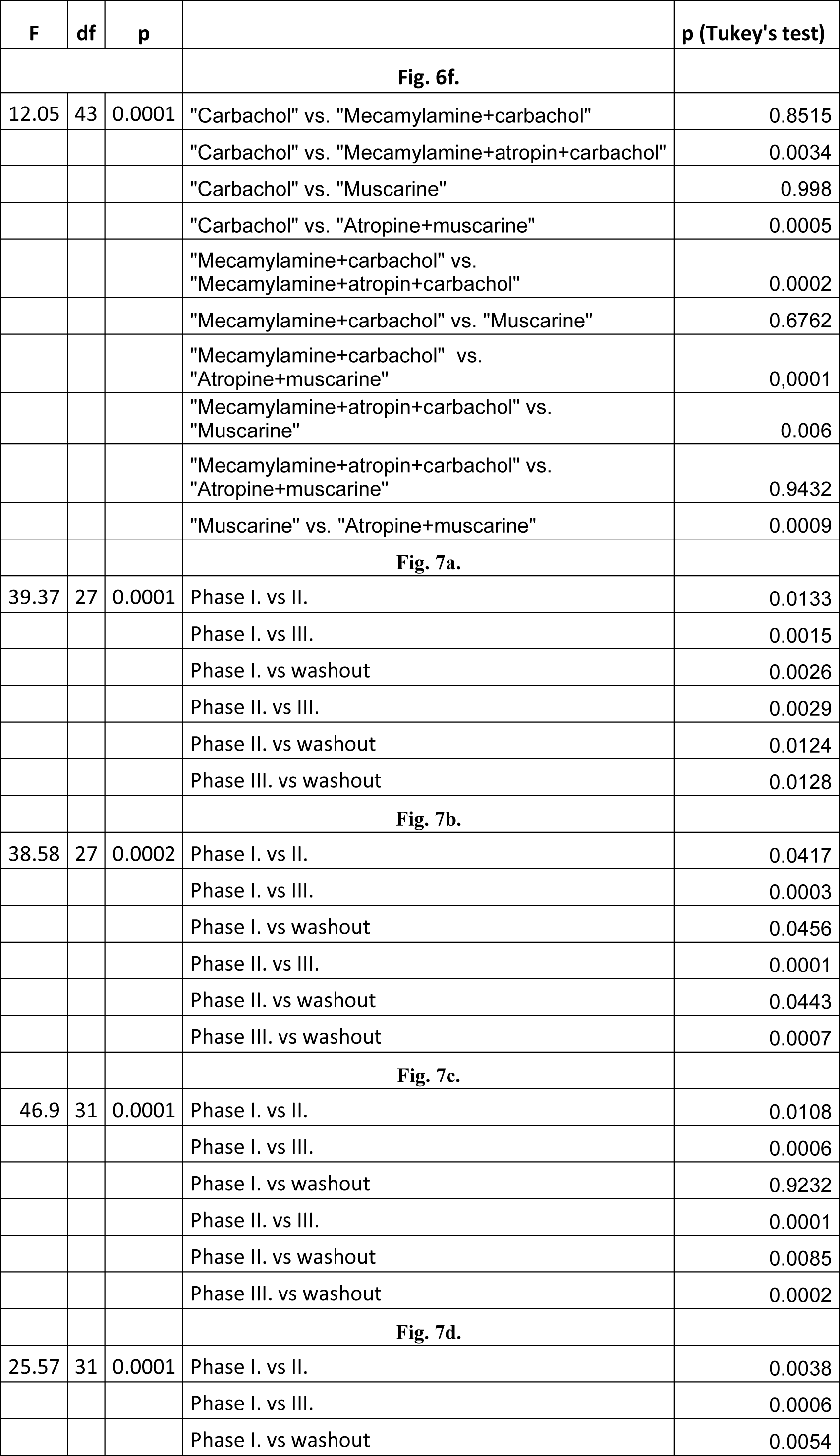
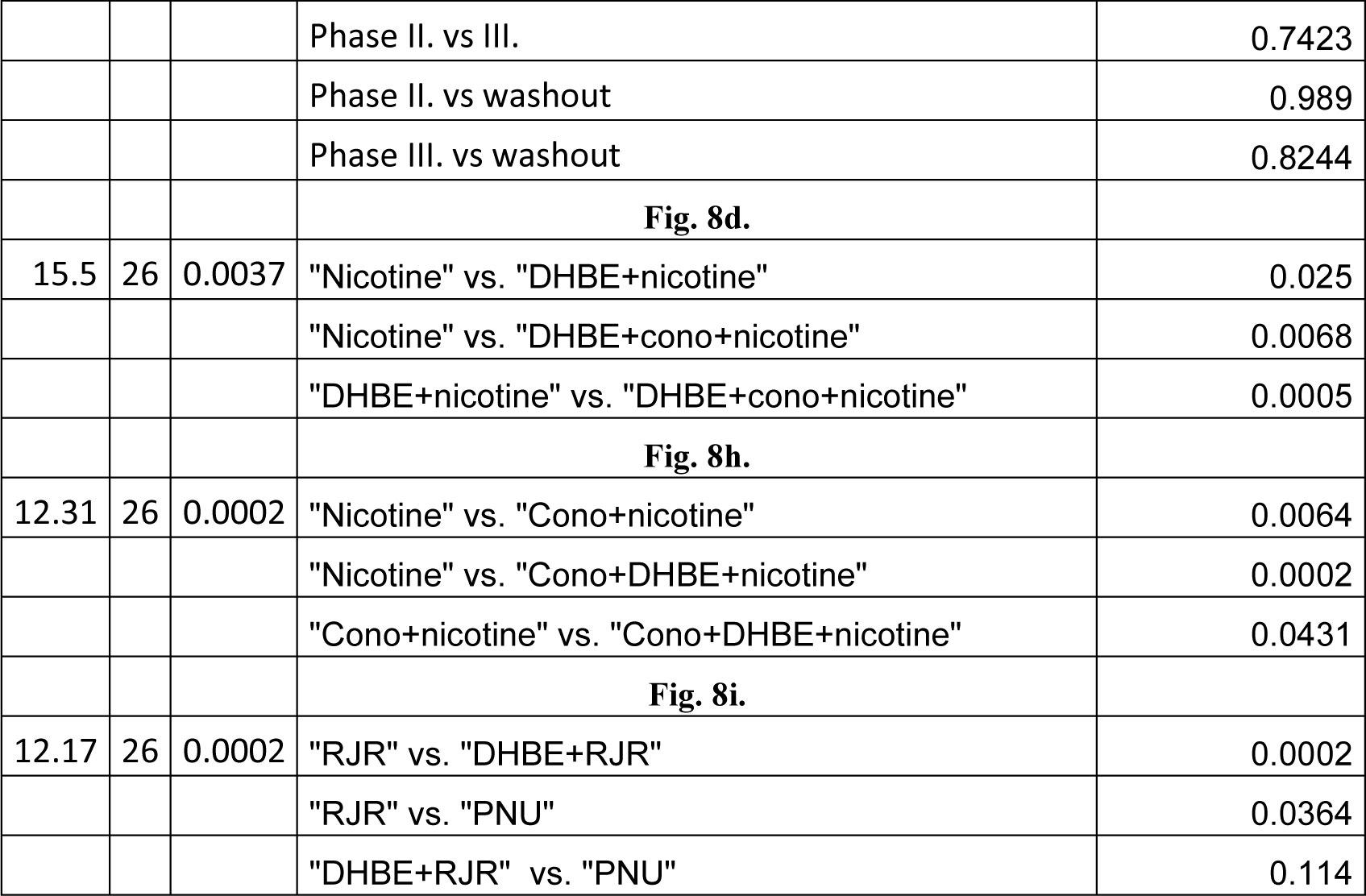
One-way ANOVA of % changes of firing rates (forFig. 6), % changes of frequencies of mPSCs (for Fig. 7), and amplitudes of the inward currents (for Fig.8)

| F | df | p |  | p (Tukey's test) |
| --- | --- | --- | --- | --- |
| **Fig. 6f.** |  |  |  |  |
| 12.05 | 43 | 0.0001 | "Carbachol" vs. "Mecamylamine+carbachol" | 0.8515 |
|  |  |  | "Carbachol" vs. "Mecamylamine+atropin+carbachol" | 0.0034 |
|  |  |  | "Carbachol" vs. "Muscarine" | 0.998 |
|  |  |  | "Carbachol" vs. "Atropine+muscarine" | 0.0005 |
|  |  |  | "Mecamylamine+carbachol" vs.<br>"Mecamylamine+atropin+carbachol" | 0.0002 |
|  |  |  | "Mecamylamine+carbachol" vs. "Muscarine" | 0.6762 |
|  |  |  | "Mecamylamine+carbachol" vs.<br>"Atropine+muscarine" | 0.0001 |
|  |  |  | "Mecamylamine+atropin+carbachol" vs.<br>"Muscarine" | 0.006 |
|  |  |  | "Mecamylamine+atropin+carbachol" vs.<br>"Atropine+muscarine" | 0.9432 |
|  |  |  | "Muscarine" vs. "Atropine+muscarine" | 0.0009 |
| **Fig. 7a.** |  |  |  |  |
| 39.37 | 27 | 0.0001 | Phase I. vs II. | 0.0133 |
|  |  |  | Phase I. vs III. | 0.0015 |
|  |  |  | Phase I. vs washout | 0.0026 |
|  |  |  | Phase II. vs III. | 0.0029 |
|  |  |  | Phase II. vs washout | 0.0124 |
|  |  |  | Phase III. vs washout | 0.0128 |
| **Fig. 7b.** |  |  |  |  |
| 38.58 | 27 | 0.0002 | Phase I. vs II. | 0.0417 |
|  |  |  | Phase I. vs III. | 0.0003 |
|  |  |  | Phase I. vs washout | 0.0456 |
|  |  |  | Phase II. vs III. | 0.0001 |
|  |  |  | Phase II. vs washout | 0.0443 |
|  |  |  | Phase III. vs washout | 0.0007 |
| **Fig. 7c.** |  |  |  |  |
| 46.9 | 31 | 0.0001 | Phase I. vs II. | 0.0108 |
|  |  |  | Phase I. vs III. | 0.0006 |
|  |  |  | Phase I. vs washout | 0.9232 |
|  |  |  | Phase II. vs III. | 0.0001 |
|  |  |  | Phase II. vs washout | 0.0085 |
|  |  |  | Phase III. vs washout | 0.0002 |
| **Fig. 7d.** |  |  |  |  |
| 25.57 | 31 | 0.0001 | Phase I. vs II. | 0.0038 |
|  |  |  | Phase I. vs III. | 0.0006 |
|  |  |  | Phase I. vs washout | 0.0054 |
|  |  |  | Phase II. vs III. | 0.7423 |
|  |  |  | Phase II. vs washout | 0.989 |
|  |  |  | Phase III. vs washout | 0.8244 |
|  |  |  | **Fig. 8d.** |  |
| 15.5 | 26 | 0.0037 | "Nicotine" vs. "DHBE+nicotine" | 0.025 |
|  |  |  | "Nicotine" vs. "DHBE+cono+nicotine" | 0.0068 |
|  |  |  | "DHBE+nicotine" vs. "DHBE+cono+nicotine" | 0.0005 |
|  |  |  | **Fig. 8h.** |  |
| 12.31 | 26 | 0.0002 | "Nicotine" vs. "Cono+nicotine" | 0.0064 |
|  |  |  | "Nicotine" vs. "Cono+DHBE+nicotine" | 0.0002 |
|  |  |  | "Cono+nicotine" vs. "Cono+DHBE+nicotine" | 0.0431 |
|  |  |  | **Fig. 8i.** |  |
| 12.17 | 26 | 0.0002 | "RJR" vs. "DHBE+RJR" | 0.0002 |
|  |  |  | "RJR" vs. "PNU" | 0.0364 |
|  |  |  | "DHBE+RJR" vs. "PNU" | 0.114 |

**Table 3.** Two-tailed Student’s t-test of % changes of firing rates (for Fig. 7 and 10), % changes of frequencies of mPSCs (for Fig. 6), and amplitudes of inward currents (for Fig. 8)

**Name of Figure**
| **Figure 6f** | **Ctrl PSC frequency (Hz)** | **Frequency of PSCs after treatment (% of ctrl)** | **N/n** | **df** | **t** | **p** |
| --- | --- | --- | --- | --- | --- | --- |
| Carbachol | 0.39 ± 0.10 | 65.7 ± 6.009 | 3/9 | 8 | 5.713 | 0.0004 |
| Mecamylamine + carbachol | 0.48 ± 0.2 | 57.2 ± 6.878 | 2/8 | 7 | 6.215 | 0.0004 |
| Mecamylamine + atropine + carbachol | 0.85 ± 0.13 | 97.2 ± 3.411 | 4/9 | 8 | 0.8144 | 0.439 |
| Muscarine | 0.97 ± 0.18 | 68.1 ± 6.67 | 4/10 | 9 | 4.767 | 0.001 |
| Atropine + muscarine | 0.91 ± 0.24 | 103.5 ± 5.116 | 4/8 | 7 | 0.6841 | 0.5159 |

| **Figure 7a** | **Ctrl firing rate (Hz)** | **Firing rate after treatment (% of ctrl)** | **N/n** | **df** | **t** | **p** |
| --- | --- | --- | --- | --- | --- | --- |
| ACh | 0.88 ± 0.25 |  | 3/7 | 6 |  |  |
| Phase I. |  | 409.3 ± 49.44 |  |  | 6.256 | 0.008 |
| Phase II. |  | 285.3 ± 39.91 |  |  | 4.642 | 0.0035 |
| Phase III. |  | 44.6 ± 8.93 |  |  | 6.207 | 0.008 |
| Washout |  | 98.7 ± 10.18 |  |  | 0.126 | 0.9036 |
| **Figure 7b** | **Ctrl firing rate (Hz)** | **Firing rate after treatment (% of ctrl)** | **N/n** | **df** | **t** | **p** |
| Carbachol | 1.0 ± 0.35 |  | 3/7 | 6 |  |  |
| Phase I. |  | 186.9 ± 20.47 |  |  | 4.242 | 0.0054 |
| Phase II. |  | 140.4 ± 9.23 |  |  | 4.38 | 0.0047 |
| Phase III. |  | 28.6 ± 7.486 |  |  | 9.542 | 0.0001 |
| Washout |  | 105.1 ± 6.27 |  |  | 0.8193 | 0.4439 |
| **Figure 7c** | **Ctrl firing rate (Hz)** | **Firing rate after treatment (% of ctrl)** | **N/n** | **df** | **t** | **p** |
| Mecamylamine + carbachol | 1.6 ± 0.7 |  | 3/8 | 7 |  |  |
| Phase I. |  | 99.4 ± 3.311 |  |  | 0.1888 | 0.8556 |
| Phase II. |  | 156.9 ± 12.16 |  |  | 4.675 | 0.0023 |
| Phase III. |  | 37.7 ± 7.67 |  |  | 8.115 | 0.0001 |
| Washout |  | 103.4 ± 5.29 |  |  | 0.6378 | 0.5439 |
| **Figure 7d** | **Ctrl firing rate (Hz)** | **Firing rate after treatment (% of ctrl)** | **N/n** | **df** | **t** | **p** |
| Atropine + carbachol | 1.6 ± 0.21 |  | 4/8 | 7 |  |  |
| Phase I. |  | 179.3 ± 12.6 |  |  | 6.289 | 0.0004 |
| Phase II. |  | 104.5 ± 4.95 |  |  | 0.9091 | 0.3935 |
| Phase III. |  | 94.9 ± 6.485 |  |  | 0.7903 | 0.4553 |
| Washout |  | 102.9 ± 3.912 |  |  | 0.735 | 0.4862 |

| **Figure 8d** | **Current amplitude (pA)** | **N/n** | **df** | **t** | **p** |
| --- | --- | --- | --- | --- | --- |
| Nicotine | -41.5 ± 9.05 | 3/9 | 8 | 4.59 | 0.0018 |
| DHBE+nicotine | -18.4 ± 3.41 | 3/9 | 8 | 5.39 | 0.0007 |
| DHBE+conotoxin+nicotine | -9.2 ± 2.36 | 3/9 | 8 | 3.909 | 0.0045 |
| **Figure 8h** | **Current amplitude (pA)** | **N/n** | **df** | **t** | **p** |
| Nicotine | -42.9 ± 7.89 | 4/9 | 8 | 5.444 | 0.0006 |
| Conotoxin+nicotine | -19.2 ± 2.83 | 4/9 | 8 | 6.771 | 0.0001 |
| Conotoxin+DHBE+nicotine | -9.2 ± 1.74 | 4/9 | 8 | 5.281 | 0.0007 |
| **Figure 8l** | **Current amplitude (pA)** | **N/n** | **df** | **t** | **p** |
| RJR | -22.9 ± 4.15 | 4/11 | 10 | 5.512 | 0.0003 |
| RJR+DHBE | -1.2 ± 1.23 | 3/8 | 7 | 0.9918 | 0.3543 |
| PNU | -11.2 ± 1.88 | 3/8 | 7 | 5.93 | 0.0006 |

| **Figure 10a** | **Ctrl firing rate (Hz)** | **Firing rate after treatment (% of ctrl)** | **N/n** | **df** | **t** | **p** |
| --- | --- | --- | --- | --- | --- | --- |
| Muscarine | 1.35 ± 0.269 |  |  |  |  |  |
| Phase I. |  | 147.6 ± 10.28 | 3/12 | 11 | 4.629 | 0.0007 |
| Phase II. |  | 54.5 ± 6.9 | 3/12 | 11 | 6.593 | 0.0001 |
| Phase I. vs Phase II. (paired t-test) |  |  | 3/12 | 11 | 11.48 | 0.0001 |
| **Figure 10b** | **Ctrl firing rate (Hz)** | **Firing rate after treatment (% of ctrl)** | **N/n** | **df** | **t** | **p** |
| Tropi+muscarine | 2.4 ± 0.58 |  |  |  |  |  |
| Phase I. |  | 152 ± 14.8 | 5/8 | 7 | 3.513 | 0.0098 |
| Phase II. |  | 78.3 ± 5.53 | 5/8 | 7 | 3.932 | 0.0057 |
| Phase I. vs Phase II. (paired t-test) |  |  | 5/8 | 7 | 6.125 | 0.0005 |
| **Figure 10c** | **Ctrl firing rate (Hz)** | **Firing rate after treatment (% of ctrl)** | **N/n** | **df** | **t** | **p** |
| Methoct+muscarine | 2.4 ± 0.38 |  |  |  |  |  |
| Phase I. |  | 145.6 ± 6.59 | 4/9 | 8 | 6.911 | 0.0001 |
| Phase II. |  | 80.8 ± 3.45 | 4/9 | 8 | 5.576 | 0.0005 |
| Phase I. vs Phase II. (paired t-test) |  |  | 4/9 | 8 | 8.872 | 0.0001 |
| **Figure 10d** | **Ctrl firing rate (Hz)** | **Firing rate after treatment (% of ctrl)** | **N/n** | **df** | **t** | **p** |
| Methoct+tropi+muscarine | 1.8 ± 0.2 |  |  |  |  |  |
| Phase I. |  | 150.8 ± 12.61 | 3/8 | 7 | 4.024 | 0.005 |
| Phase II. |  | 122.4 ± 8.76 | 3/8 | 7 | 2.555 | 0.0378 |
| Phase I. vs Phase II. (paired t-test) |  |  | 3/8 | 7 | 2.292 | 0.0557 |
| **Figure 10e** | **Ctrl firing rate (Hz)** | **Firing rate after treatment (% of ctrl)** | **N/n** | **df** | **t** | **p** |
| Piren+methoct+tropi+muscarine | 0.9 ± 0.18 |  |  |  |  |  |
| Phase I. |  | 118.9 ± 4.5 | 3/8 | 7 | 4.189 | 0.0041 |
| Phase II. |  | 102.4 ± 6.42 | 3/8 | 7 | 0.3696 | 0.7226 |
| Phase I. vs Phase II. (paired t-test) |  |  | 3/8 | 7 | 2.649 | 0.033 |
| **Figure 10f** | **Ctrl firing rate (Hz)** | **Firing rate after treatment (% of ctrl)** | **N/n** | **df** | **t** | **p** |
| Dari+methoct+tropi+muscarine | 1.2 ± 0.16 |  |  |  |  |  |
| Phase I. |  | 119.4 ± 4.18 | 4/9 | 8 | 4.648 | 0.0016 |
| Phase II. |  | 124.0 ± 9.71 | 4/9 | 8 | 2.471 | 0.0386 |
| Phase I. vs Phase II. (paired t-test) |  |  | 4/9 | 8 | 0.4587 | 0.6587 |
| **Figure 10g** | **Ctrl firing rate (Hz)** | **Firing rate after treatment (% of ctrl)** | **N/n** | **df** | **t** | **p** |
| Dari+piren+methoct+tropi+muscarine | 1.2 ± 0.61 |  |  |  |  |  |
| Phase I. |  | 105.3 ± 8.31 | 3/10 | 9 | 0.6375 | 0.5397 |
| Phase II. |  | 103.2 ± 7.89 | 3/10 | 9 | 0.4055 | 0.6945 |
| Phase I. vs Phase II. (paired t-test) |  |  | 3/10 | 9 | 0.2339 | 0.8203 |

**Table 4.** mPSC frequency responses upon blocking inhibitory or stimulatory muscarinic receptors and adding muscarine in the absence or presence of THL (Student’s t-test and ANOVA;Fig. 9)

| Before muscarine | Inhibition of | Control frequency (Hz) | Frequency after treatments (Hz) | Frequency after treatments (% of control) | n/N | Student's t-test |  |  |
| --- | --- | --- | --- | --- | --- | --- | --- | --- |
|  |  |  |  |  |  | df | t | p |
| Without THL | Inhibitory receptors | 1.8 ± 0.10 | 2.3 ± 0.23 | 127.3 ± 8.50 | 10/3 | 9 | 3.213 | 0.0106* |
|  | Stimulatory receptors | 1.6 ± 0.12 | 1.3 ± 0.11 | 80.7 ± 2.67 | 10/4 | 9 | 7.226 | 0.0001* |
| With THL | Inhibitory receptors | 1.3 ± 0.22 | 1.4 ± 0.24 | 102.2 ± 4.1 | 10/3 | 9 | 0.5404 | 0.602 |
|  | Stimulatory receptors | 1.3 ± 0.20 | 1.3 ± 0.20 | 102.3 ± 5.1 | 10/4 | 9 | 0.4538 | 0.6607 |
|  |  |  |  |  |  | ANOVA |  |  |
|  |  |  |  |  |  | df | F | p |
|  |  |  |  |  |  | 39 | 11.94 | 0.0001* |
| After muscarine | Inhibition of | Control frequency (Hz) | Frequency after treatments (Hz) | Frequency after treatments (% of control) | n/N | Student's t-test |  |  |
|  |  |  |  |  |  | df | t | p |
| Without THL | Inhibitory receptors | 2.5 ± 0.48 | 3.1 ± 0.39 | 139.1 ± 14.75 | 10/3 | 9 | 2.651 | 0.0264* |
|  | Stimulatory receptors | 1.3 ± 0.07 | 0.9 ± 0.08 | 73.4 ± 5.09 | 10/4 | 9 | 5.223 | 0.0005* |
| With THL | Inhibitory receptors | 1.4 ± 0.15 | 1.4 ± 0.14 | 106.5 ± 4.17 | 10/3 | 9 | 1.56 | 0.1532 |
|  | Stimulatory receptors | 1.6 ± 0.31 | 1.6 ± 0.26 | 106.4 ± 4.97 | 10/4 | 9 | 1.289 | 0.2295 |
|  |  |  |  |  |  | ANOVA |  |  |
|  |  |  |  |  |  | df | F | p |
|  |  |  |  |  |  | 39 | 10.08 | 0.0001* |

n = number of recorded neurons; N=number of animals
df = degree of freedom
t = "t" values of Student's t-test
F = "F" value of ANOVA
p = "p" probability values of % data
\* = significant change

## Experimental design and statistical analysis

Gonadectomy (GDX) was performed before CNO treatment to raise the frequency of LH pulses in the absence of the negative feedback effect of the gonadal steroids. Using the Chat-Gq (hM3Dq) DREADD mouse model, cholinergic neurons were selectively activated by administering CNO (5 mg/kg body weight; n = 7) or vehicle DMSO (n = 6) orally to GDX mice. The LH levels were monitored using an ultrasensitive ELISA assay (Steyn et al., 2013) with tail-tip bleeding samples taken at 3-minute intervals over a period of 3 hours for both experimental groups.

Based on the 3DISCO clearing technique, a high-throughput double immunofluorescent study was used to find a cholinergic network that targets the GnRH neurons. We used 1-mm thick brain slices of adult GnRH-GPF male mice (n = 6) cut in the para-medial sagittal plane bilaterally to maintain the structural integrity of the region of interest for the purpose of image reconstruction. Using confocal microscopy, optical sections were obtained, and image stacks were reconstructed. The distribution of VAChT immunoreactive inputs (juxtapositions) on GnRH dendrites and soma was analyzed, along with the innervation ratio of GnRH perikarya.

The cholinergic innervation of GnRH neurons was characterized at the ultrastructural level through the distinction of chromogens used in double labeling of VAChT (nickel-DAB) and GnRH (silver-intensified colloidal gold particles) immunoreactive profiles, enabling clear identification of juxtapositions and synapses at the ultrastructural level.

To confirm synaptic innervation of GnRH-cre neurons by central cholinergic neurons and identify the subgroups of ChAT-IR neurons with that potential, the septo-preoptic area populated by GnRH-Cre neurons was infected in two steps with genetically altered helper and rabies viruses. Helper virus-infected GnRH-Cre neurons expressing TVA receptor and G proteins became targets of the modified, envelope protein A (EnvA)-coated, G-deleted rabies virus (RVdG), which then labeled the monosynaptically connected afferent neurons with their fluorescent transgene. The retrograde tracing was combined with fluorescent ChAT-immunocytochemistry to identify the cholinergic phenotype of the afferent neurons.

Using TaqMan PCR, the relative gene expression (mRNA) levels of cholinergic receptor subunits were calculated in pooled GnRH cytoplasmic samples (n = 7) to determine whether GnRH neurons are direct targets of cholinergic neurotransmission.

Inward membrane currents and postsynaptic currents (PSCs) in GnRH neurons were recorded in whole-cell voltage clamp mode at −70 mV holding potential. Various selective and non-selective AChR agonists (ACh, carbachol, nicotine, muscarine, etc.) were applied in a single bolus into the measurement chamber directly onto the acute brain slice, and changes in the membrane current or in the frequency of PSCs were measured. These measurements were repeated in the presence of various cocktails of selective and non-selective AChR antagonists (mecamylamine, atropine, DHβE, ω-conotoxin, darifenacin, pirenzepine, etc.) to demonstrate the role of the diverse nicotinic and/or muscarinic AChRs in the observed effects. Statistical analyses are shown in the corresponding statistical tables.

Changes in the firing rate of GnRH neurons resulting from activation of AChRs were proven in whole-cell current clamp mode at 0 pA holding current. AChRs agonists (ACh, carbachol, or muscarine) were applied in a single bolus, and then the change in firing rate was examined. These measurements were then repeated in the presence of various cocktails of AChR antagonists. Detailed statistical analyses are given in the statistical tables.

Involvement of the endogenous ACh release from the axon terminals contacting GnRH neurons was investigated by optogenetic examinations in brain slices from 3x-transgenic mice, expressing channelrhodopsin in the boutons containing ACh. LED illumination of the acute brain slices at 470 nm and 5 Hz was used to release ACh from these boutons, and changes in the firing rate or frequency of PSCs were recorded. The measurements were repeated in the presence of mecamylamine/atropine. Statistical analyses can be found in the statistical tables.

Frequency-dependent GABA *versus* ACh synaptic release to GnRH neurons from the channelrhodopsin-expressing ACh-containing axon terminals was revealed in brain slices by optogenetics from triple-transgenic mice. LED illumination of two frequencies (0.2 Hz *versus* 5 Hz) was used to evoke GABAergic and/or cholinergic PSCs. Then the measurements were repeated in the presence of the GABA_A_-R antagonists picrotoxin and mecamylamine/atropine, respectively.

## Results

### Pharmacogenetic activation of the central cholinergic system *in vivo* increases secretion of luteinizing hormone (LH) in orchidectomized mice

To study the effect of increased brain acetylcholine transmission upon secretion of luteinizing hormone (LH), a pharmacogenetic approach was used in transgenic *Chat-Cre-Gq DREADD* mice. Gonadal hormones are known to suppress the HPG axis via negative feedback; therefore, orchidectomized (ORX) animals were used that were accustomed to chronic handling prior to the experiment. The animal groups were composed of vehicle (DMSO) and clozapine-N-oxide (CNO)-administered mice. Blood samples were collected every 3 minutes over a 3 hour period and their luteinizing hormone (LH) content was measured using an ultrasensitive ELISA technique (Steyn et al., 2013). We found that in the control group, DMSO administration had no influence upon LH secretion; the corresponding pre-treatment (basal LH: 4.92 ± 0.55 ng/ml; mean LH: 6.03 ± 0.58 ng/ml) and post-treatment (basal LH: 4.99 ± 0.59 ng/ml; mean LH: 6.17 ± 0.64 ng/ml) values didn’t show significant differences (**Fig. 1 a-c**). In contrast, administration of CNO evoked a significant rise in both the basal (pre-treatment basal LH: 9.20±1.53 ng/ml vs. post-treatment basal LH: 18.32 ± 2.21 ng/ml, paired t-test, p = 0.0104) and mean (pre-treatment mean LH: 10.97.21±1.71 ng/ml vs. post-treatment mean LH: 23.18 ± 3.51 ng/ml, paired t-test, p = 0.0255) LH values. (**Fig. 1 d-f**). The LH values peaked 30 minutes after the initiation of CNO treatment.

**Fig. 1.**
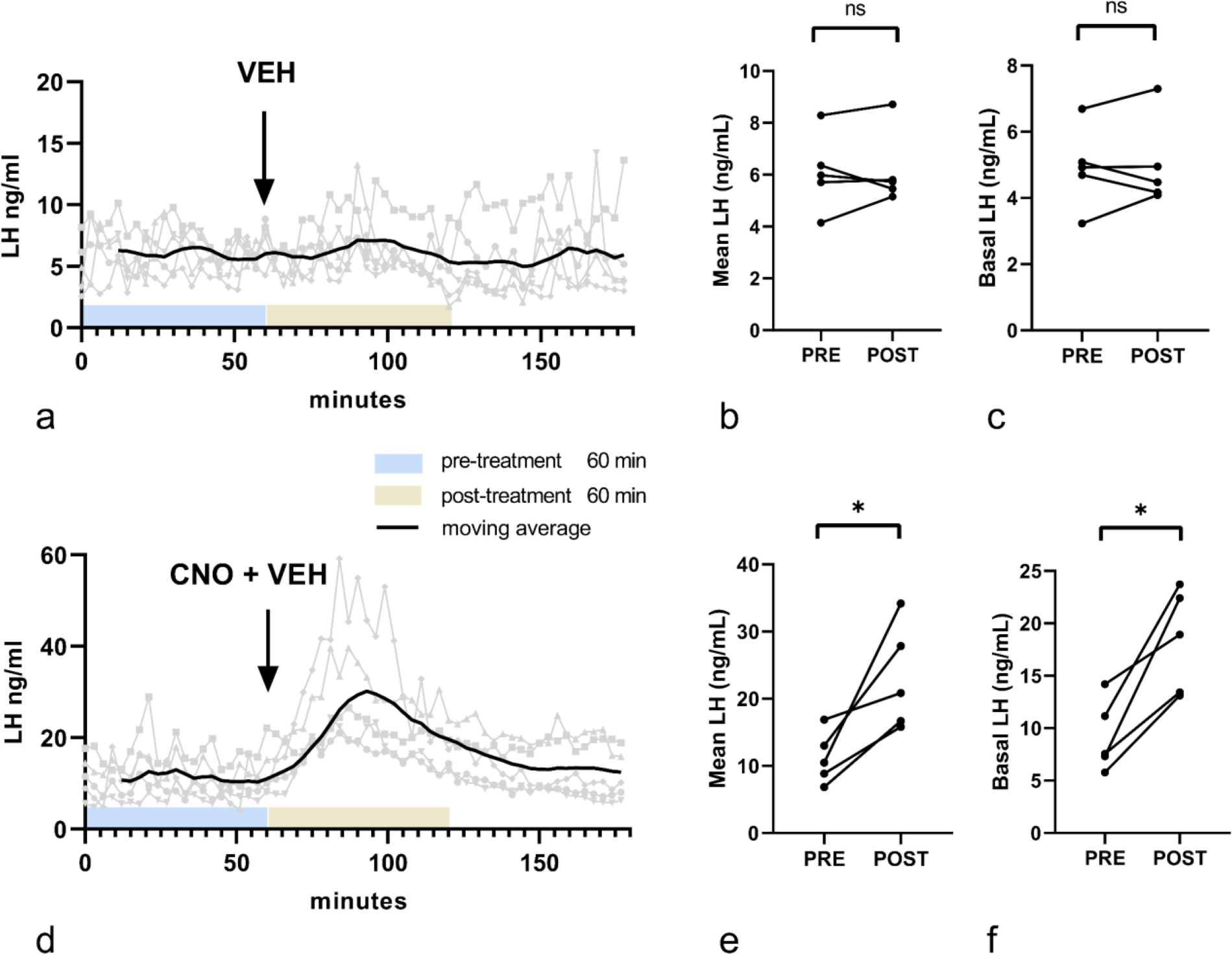
Effect of clozapine N-oxide (CNO) treatment on serum LH level in orchidectomized Chat-Cre-Gq DREADD mice. Blood samples were collected every 3 minutes over a 3-hour period. The LH concentration was determined using an ultrasensitive ELISA technique. The control (**a-c**) and CNO-treated (**d-f**) mice received vehicle or CNO, respectively, per os at the 60^th^ minute from the onset of sampling. Graphs **a** and **d** show mean LH levels per sampling time revealed by ELISA. Comparing individual animals, the mean (**b**) and basal (**c**) LH serum levels remained unchanged within a 60-minute pre- and post-treatment period in the vehicle-treated control mice. In contrast, CNO treatment significantly increased both mean (**e)** and basal (**f**) serum LH concentration in the 60-minute post-treatment period (paired t-test; *p < 0.05).

**Fig. 2.**
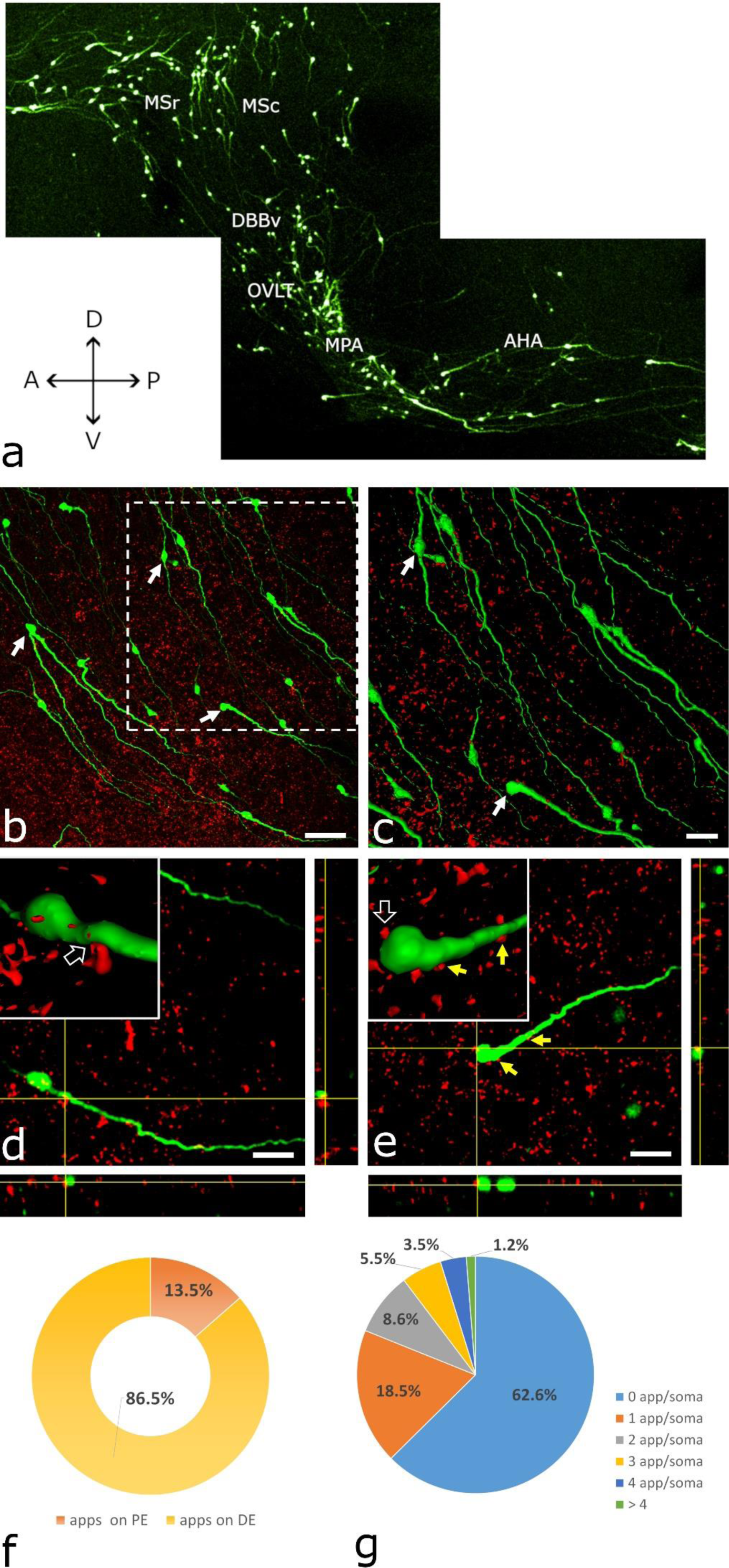
GnRH neurons receive cholinergic inputs as revealed by double immunostaining and the 3DISCO method. **a.** Distribution of GnRH-GFP-IR elements revealed by 3DISCO. A thick brain slice (1 mm) cut in the paramedian sagittal plane. It represents the scanning areas used for image acquisition via confocal microscopy. **b**. A stack of confocal image planes projected in the z direction shows many GnRH neurons (green channel, arrows) and VAChT-immunoreactive (IR) axon varicosities (red channel). **c**. A part of the image stack shown by the dashed square in **Fig. 2b** was reconstructed as a 3D surface model using the ImageJ/Fiji software. Note the overlap of the two immunolabeled systems. **c.** Axo-dendritic contact between a VAChT-IR axon and a GnRH neuron is shown in both the 3D surface model (inset, arrow) and orthogonal views of stacks (yellow crosshairs). **e.** The inset shows an axo-somatic juxtaposition (empty arrow) in a 3D image. The same communication site is confirmed in orthogonal views (yellow crosshairs). Two other cholinergic axon varicosities (yellow arrows) juxtaposed to the same GnRH neuron indicate the existence of multiple contact sites. **f.** The cholinergic axons communicate with GnRH dendrites (86.5%); the perikarya are less often targeted (13.5%). Apps on PE: appositions on perikarya; Apps on DE: appositions on dendrites. **g.** About one third (37,4 %) of the analyzed GnRH perikarya received VAChT-IR inputs. The inner pie diagram depicts the percentage distribution of VAChT-IR axon beads juxtaposed to the same GnRH perikaryon. App/soma: apposition per soma. Scale bars: a: 50 µm; b: 25 µm; c and d: 20 µm. MSr: medial septum, rostral part; MSc: medial septum, caudal part; DBBv: diagonal band of Broca, vertical part; OVLT organum vasculosum of the lamina terminalis; MPA: medial preoptic area and AHA: anterior hypothalamic area.

### Networking of central cholinergic neurons with GnRH neurons

#### GnRH neurons receive cholinergic innervation

For structural evaluation and quantitative analysis of the putative interneuronal communication of the central cholinergic and GnRH systems, double immunofluorescence labeling combined with the 3DISCO method was used on paramedian sagittal slices of the GnRH-GFP transgenic mouse brain that contained the distribution sites of GnRH neurons in the medial septum, the vertical limb of the diagonal band of Broca, the region of the organum vasculosum laminae terminalis, the medial preoptic area, and the anterior hypothalamic area (**Fig. 2a**). The cholinergic axons were identified by their vesicular acetylcholine transporter (VAChT) content, whereas GnRH-immunoreactive (IR) profiles were visualized by immunostaining of the expressed green fluorescent protein (GFP). GnRH neurons were found to overlap with VAChT-IR axons **Fig. 2b and 2c**) distributed in the sampled regions. Altogether, 4,899 optical slices (mean ± SEM: 816 ± 64 per brain) were evaluated. In each optical slice, the number of VAChT-IR axons juxtaposed to GnRH-IR perikarya and dendrites was determined. Juxtapositions were justified with high precision in the orthogonal view layout. Many VAChT-IR axon varicosities (1040 ± 96 SEM, n = 6) were identified in juxtaposition with GnRH-IR profiles (**Fig. 2d and 2e**). Most of them (86.5 ± 3.2%) targeted GnRH dendrites (**Fig. 2d**) and only a smaller fraction (13.5 ± 3.2 percent) contacted GnRH perikarya (909 ± 100.12 dendritic vs. 131 ± 23.68 somatic input, n = 6; paired t-test, p = 0.0008, two-tailed; **Fig. 2e, f**). In these samples, as an average, 37.4 ± 5.1 percent of reconstructed GnRH-IR perikarya (186 ± 20) received cholinergic inputs (68 ± 9.66 innervated vs. 118 ± 16.9 non-innervated GnRH-IR perikarya, n = 6; paired t-test, p = 0.0454, two-tailed; **Fig. 2g**).

#### Ultrastructural correlates of networking

The networking among VAChT-IR axons and GnRH neurons was also confirmed at the ultrastructural level using pre-embedding double immunocytochemistry. For the labeling of VAChT-IR axons, nickel-3,3 diaminobenzidine (Ni-DAB) chromogen was used (**Fig. 3a and 3b**), while the GnRH immunoreactive profiles were visualized by silver-intensified colloidal gold particles (**Fig. 3c and 3d**). In the residence area of hypophysiotropic GnRH neurons, the medial septum-diagonal band of Broca-OVLT continuum, VAChT-expressing axons appeared in juxtaposition to cell bodies and dendrites (**Fig. 3a**) and formed synapses (**Fig. 3b**). Based upon the different physico-chemical properties of the used chromogens, the GnRH and VAChT-IR profiles were clearly distinguishable from each other even at medium-power electron microscopic screening (**Fig. 3d**). The juxtaposition of cholinergic axon beads to dendritic and somatic domains of GnRH neurons without glial process interposition was confirmed at the ultrastructural level. Tracing the juxtaposed profiles through a series of ultrathin sections occasionally revealed the synaptic engagement of VAChT-IR, cholinergic axons with dendrites (**Fig. 3e**) and perikarya (**Fig. 3f and 3g**) of GnRH neurons, suggesting a direct regulatory influence of the cholinergic system upon the GnRH neuron assembly.

**Fig 3.**
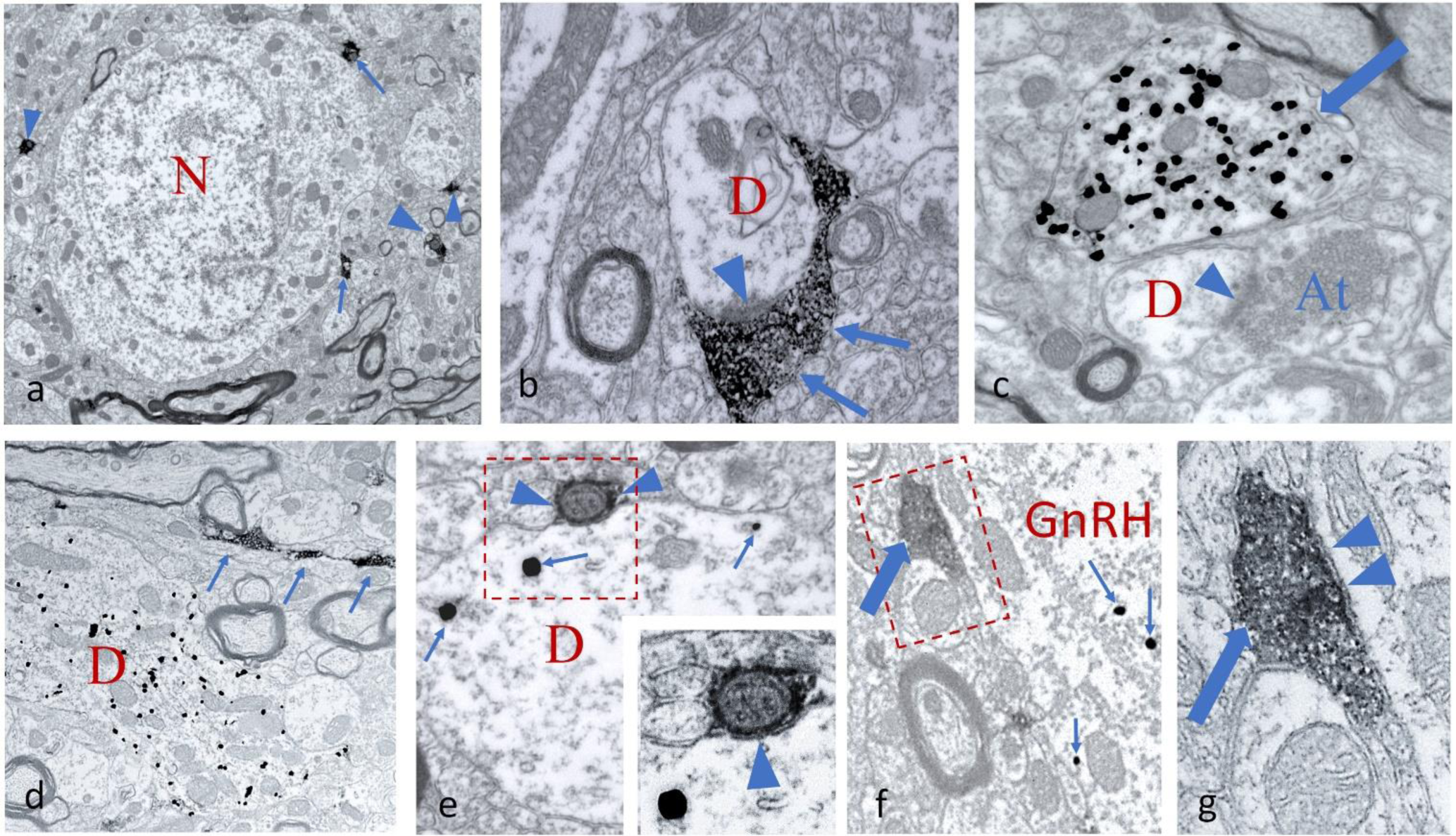
Ultrastructural detection of cholinergic axons and GnRH-immunoreactive (IR) profiles in the medial preoptic area (mPOA) **a.** Vesicular acetylcholine transporter (VAChT)-containing axons labeled by Ni-DAB chromogen are distributed in the neuropil of the mPOA, contacting cell bodies (arrows) and dendrites (arrowheads). A non-labeled neuron (N) is in the center of micrograph. **b**. A Ni-DAB chromogen-labeled cholinergic axon (arrows) synapses (arrowhead) on a cross-sectioned dendrite (D). **c**. Appearance of a cross-sectioned GnRH dendrite (arrow) filled with silver-intensified, colloidal gold particles. A nearby unlabeled dendrite (D) receives a synapsing (arrowhead) axon terminal (At). **d**. Simultaneous detection of a longitudinally sectioned GnRH dendrite (D) identified by silver-coated colloidal gold particles and a varicose, longitudinally cut VAChT-IR axon (arrows) labeled by Ni-DAB chromogen. **e**. Cross-sectioned GnRH-IR dendrite (D) communicating with a small caliber cholinergic axon (arrowheads). Arrows point to the metallic label within the GnRH profile. The enframed region is shown at higher power in the inset, and the arrowhead points to the postsynaptic membrane. **f.** Detail from a GnRH perikaryon (GnRH) that receives a cholinergic axon (thick arrow). Intensified, colloidal gold particles are labeled by small arrows. The communication site is enframed. **g**. High-power view of the enframed area in **Fig. f.** The labeled, cholinergic axon (thick arrow) filled with small-sized vesicles establishes a synapse (arrowheads) with the GnRH perikaryon. Scale bars: **a**: 2 µm; **d**: 1 µm; **b** and **c**: 500 nm; **e-g**: 250 nm.

### Identification of cholinergic neuronal source supplying innervation of GnRH neurons

To figure out which of the brain regions host cholinergic cells innervating GnRH neurons, a monosynaptic, rabies virus retrograde tract tracing technique was used (Wall et al., 2010; Roelofs et al., 2021) in combination with immunocytochemistry. GnRH-Cre neurons (**Fig. 4a and 4b**) expressing TVA receptor and G proteins became selectively infected by the modified, envelope protein A (EnvA)-coated, G-deleted rabies virus (RVdG). GnRH neurons expressed the TVA-GFP/TVA receptor protein as the precursor neurons for rabies infection. (**Fig. 4a**). No mCherry fluorescence of helper virus origin was seen in wild-type mice, showing that the viral activation was Cre-dependent. Rabies virus-labeled neurons forming the primary afferents of the GnRH-Cre cells appeared throughout the entire length of the neuroaxis, with the highest number observed in the vicinity of the starter cells. Proceeding from the preoptic area toward the brain stem, their number gradually diminished. Detection of GFP fluorescence, an indicator of rabies virus transfection, in choline acetyltransferase (Chat)-immunoreactive neurons identified the source of cholinergic afferents to GnRH neurons. These double-labeled neurons were in the medial septum (MS) and the diagonal band of Broca (DBB) (**Fig. 4c and 4d**). No primary cholinergic afferent neurons were identified in other parts of the forebrain or in the brainstem, indicating that the cholinergic regulatory input is local to the residence of GnRH neurons.

**Fig 4.**
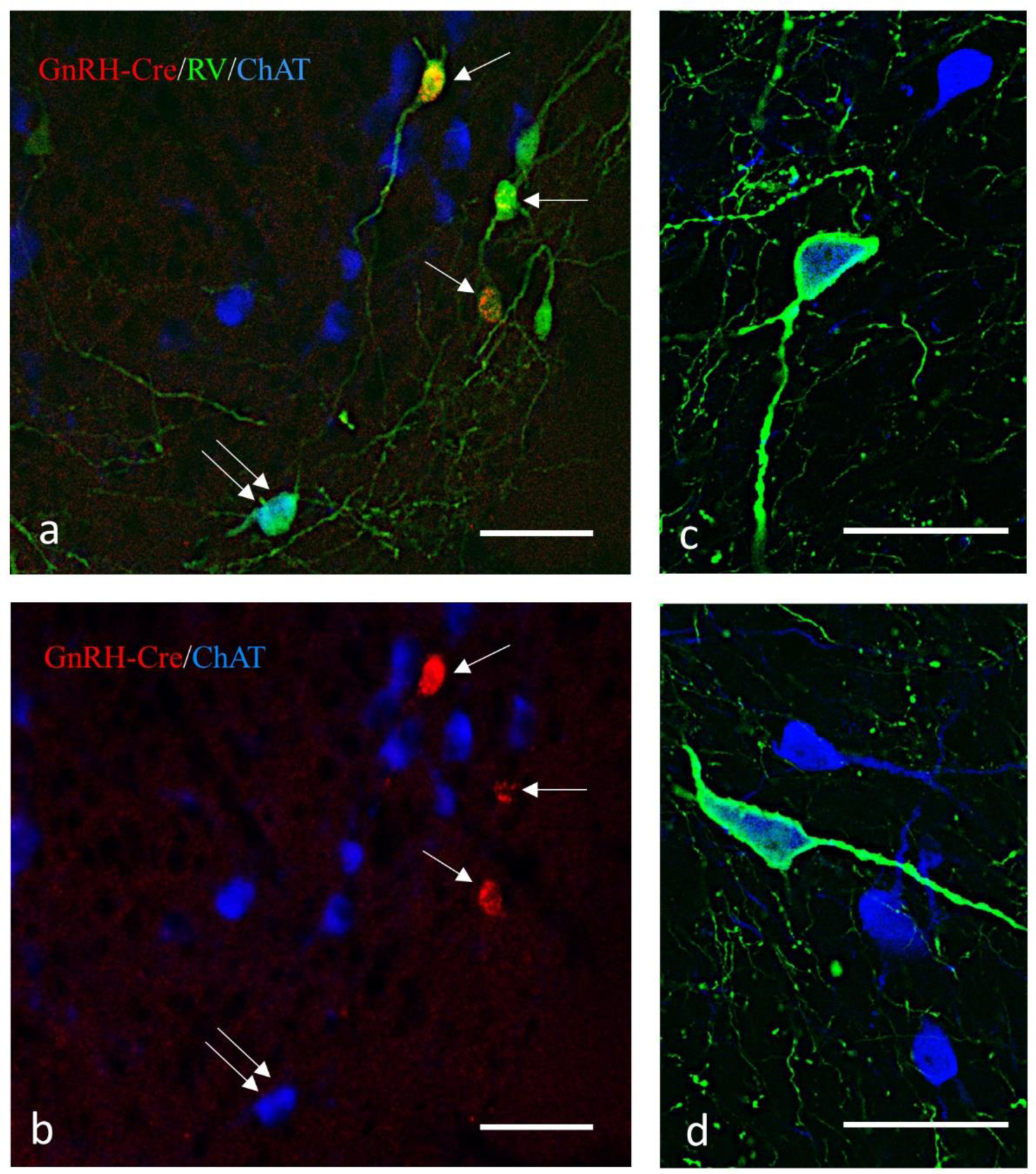
Viral tracing of cholinergic afferents of the GnRH neurons. Monosynaptic tracing of primary afferents to GnRH-Cre neurons and subsequent immunohistochemical processing of brain sections identify AAV-infected, Cre-positive neurons expressing mutated TVA and G protein fused to mCherry (in red), rabies virus-GFP infected cells (in green), and Chat-immunoreactive (IR) cells (in blue). Contrary to the mCherry signal, the green fluorescence fills both the perikarya and the processes. **a.** Starter cells (single arrows) express both mCherry and GFP, whereas the primary afferent neurons produce only GFP. Double arrows point to a cholinergic primary afferent neuron (in turquoise) having a mixture of both green and blue signals. **b.** Validation of double-labeled neurons by showing the fluorescent signals in single channels. **c and d**. Examples of Chat-IR cells labeled with the rabies GFP signal in the septo-diagonal band area of the mouse brain. Scale bar: 50 µm.

### Hypophysiotropic GnRH neurons express various nicotinic and muscarinic ACh receptors

Cytoplasmic samples (n = 70) were collected from GnRH-GFP neurons by patch pipette aspiration, sorted into separate cytoplasmic sample pools (CPs, n = 7), and examined by quantitative real-time PCR (qRT-PCR). The primer probe sets used for detection of expression of genes encoding for muscarinic and nicotinic ACh receptors, housekeeping genes (*Hprt*, *Gapdh*), and *Gnrh1* are summarized in **Table 1**. Examination of the sample groups revealed the relative mRNA expression level of genes encoding for the following nicotinic receptor subunits in GnRH neurons (mean ± SE, **Fig 5b**): alpha 3 subunit (*Chrna*3) 0.026 ± 0.006 (detected in 2 of 7 CPs), alpha 4 subunit (*Chrna4*) 0.016 ± 0.009 (2 of 7 CPs), alpha 7 subunit (*Chrna7*) 0.018 ± 0.009 (3 of 7 CPs), beta 2 subunit (*Chrnb2*) 0.015 ± 0.005 (4 of 7 CPs); beta 3 subunit (*Chrnb3*) 0.023 ± 0.11 (4 of 7 CPs), and beta 4 subunit (*Chrnb4*) 0.047 ± 0.016 (5-of 7 CPs). Regarding muscarinic receptors (M1–M5), muscarinic type 1 (*Chrm1*) 0.034 ± 0.004 (5 of 7 CPs), type 2 (*Chrm2*) 0.011 ± 0.006 (2 of 7 CPs), type 3 (*Chrm3*) 0.048 ± 0.016 (5 of 7 CPs), type 4 (*Chrm4*) 0.038 ± 0.013 (4 of 7 CPs), and type 5 (*Chrm5*) 0.037 ± 0.024 (3 of 7 CPs) receptor genes were also measured in the samples (**Fig. 5b**.) The expression of the *Gnrh* gene (30.2 ± 4.68) was confirmed in all samples (**Fig. 5a**).

**Fig 5.**
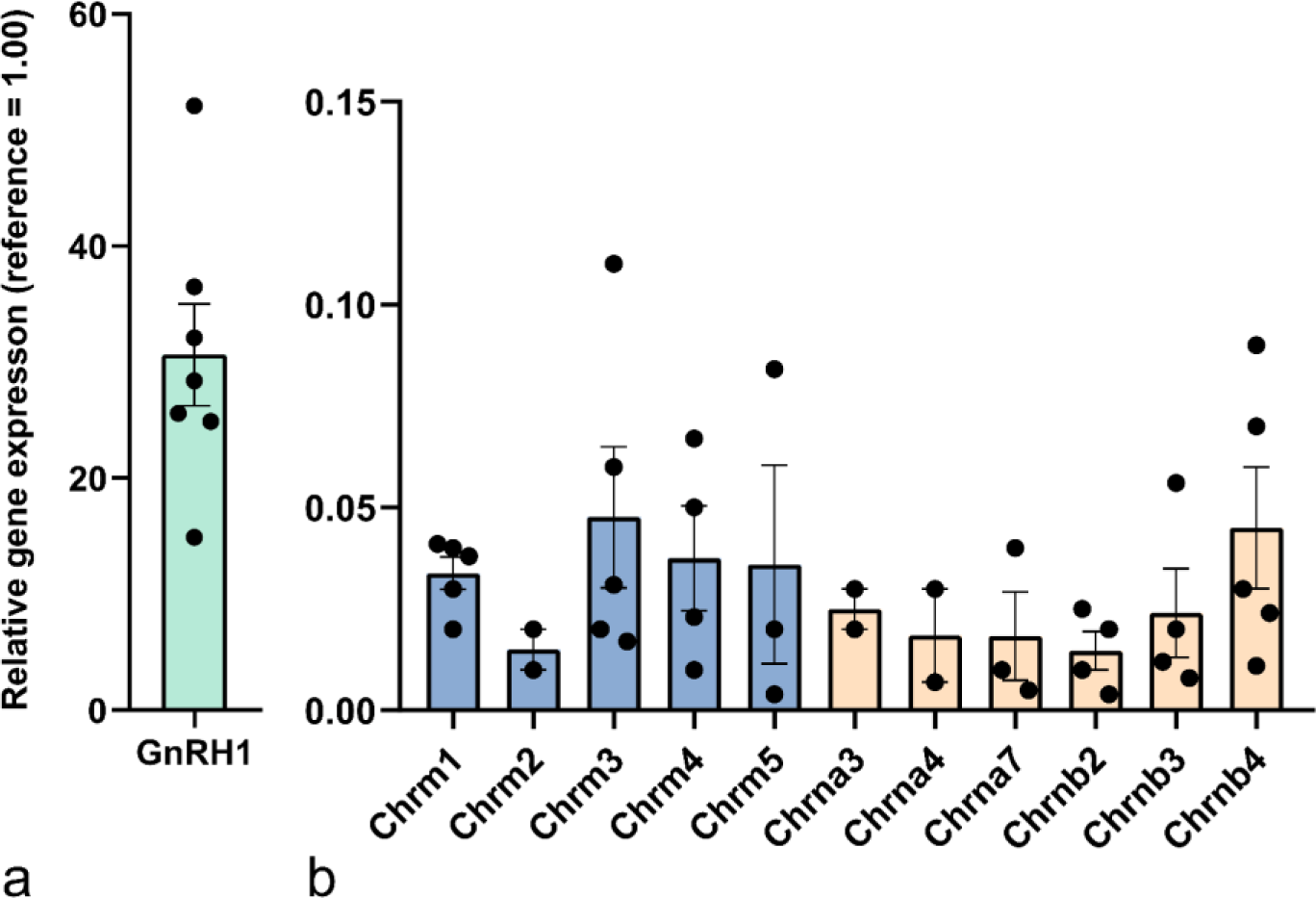
Expression of nicotinic and muscarinic ACh receptor genes in GnRH neurons. Relative expression of the Gnrh1 gene (**a**) and various nicotine and muscarine acetylcholine receptor genes (**b**) in cytoplasmic sample pools of GnRH neurons were revealed by qRT-PCR. The mean ± standard error of gene expression data is shown for each subunit as a proportion of the mean of the two reference genes.

### Cholinergic drugs change the electrophysiological activity of GnRH neurons in acute brain slice

#### Carbachol triggers ion current and modifies mPSCs of GnRH cells

We studied the effects of activation of nicotinic and muscarinic AChRs by the AChR agonist, carbachol, on the membrane current and miniature postsynaptic currents (mPSCs) in voltage-clamp mode at −70 mV. Application of carbachol in a single bolus triggered a transient inward current (amplitude = −36.9 ± 3.09 pA, Student’s t-test: N/n=3/9, p=0.0001, t=11.95, df=8; duration = 47.9 ± 12.34 s) (**Fig. 6a**). Thereafter, the frequency of mPSCs significantly decreased (65.7 ± 6.01 % of the control value, **Fig 6a**). We hypothesized that the inward current is caused by nAChR. Therefore, we next pretreated the brain slices with mecamylamine (non-selective nAChR antagonist, 10 μM), and then carbachol was added to the aCSF (**Fig. 6b**). The disappearance of the inward current confirmed that nAChR activation is involved in the effect of carbachol on membrane current. Nevertheless, the significant decrease in the frequency of the mPSCs was still observable (57.2 ± 6.88% of the control value, (**Fig. 6b**) raising the possibility that mAChRs play a role in the regulation of mPSC frequencies. Thus, next, the slices were pretreated with a cocktail of mecamylamine (10 μM) and the non-selective mAChR antagonist, atropine (10 μM), and then carbachol was added to the aCSF (**Fig. 6c**). The inward current disappeared, and the frequency of mPSCs remained unchanged in this experiment (frequency: 97.2 ± 3.41 % of the control value, **Fig 6c**).

**Fig. 6.**
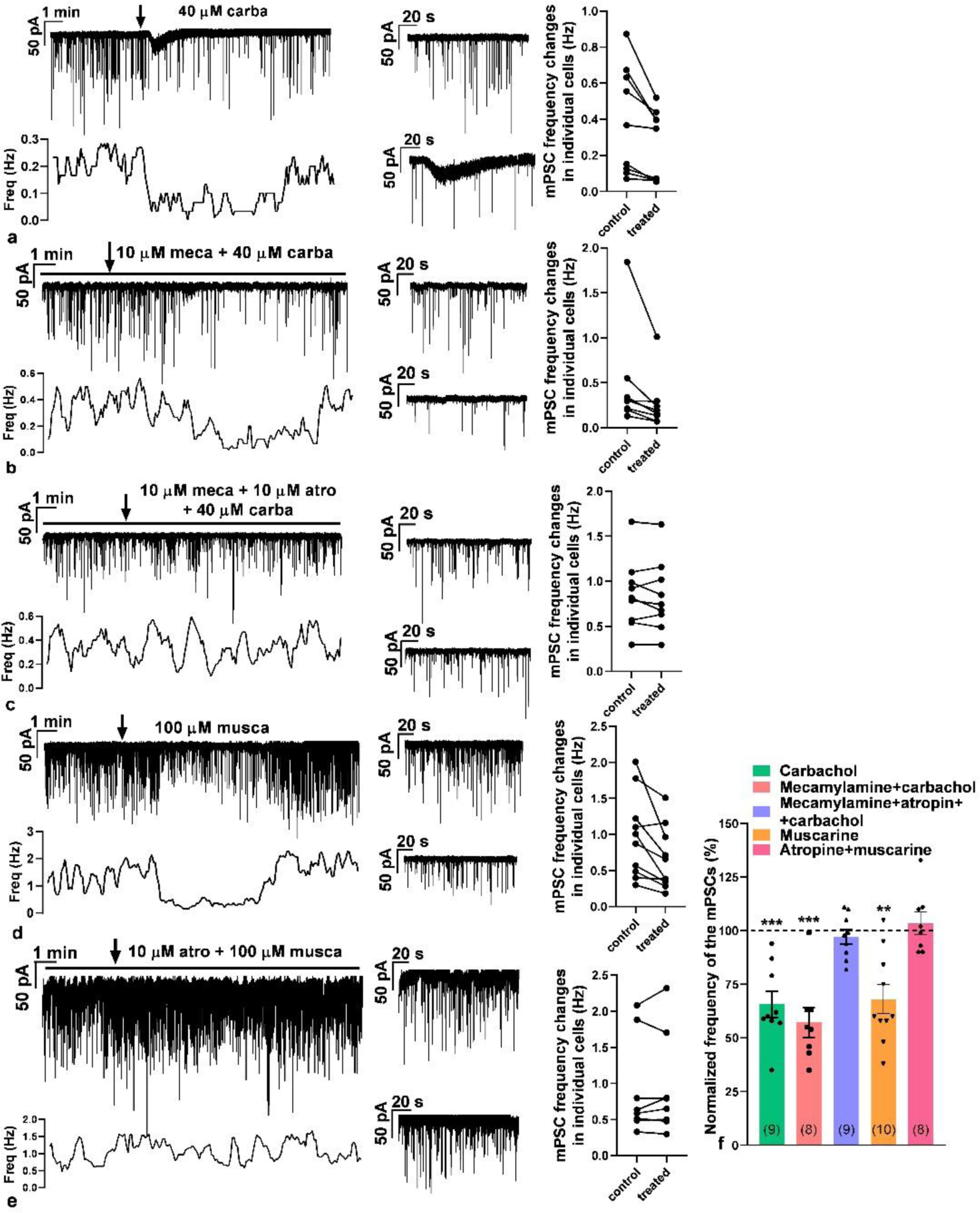
Effect of carbachol on the membrane current and mPSCs of GnRH neurons. Representative recordings illustrate that carbachol (40 μM) first evoked an inward current and then an inhibitory action on mPSCs of GnRH neurons in the presence of tetrodotoxin (660 nM). **a**. Carbachol triggered an inward current, followed by a slower inhibitory effect on the frequency of mPSCs. **b**. Mecamylamine pretreatment (10 μM) revealed that the inward current is related to nicotinic receptors. **c**. Simultaneous pretreatment with mecamylamine and atropine (10 μM) eliminated both the inward current and the inhibitory effect of carbachol. **d**. Muscarine 100 μM) evoked exclusively the inhibitory effect upon mPSCs. **e**. This inhibitory effect was totally abolished by atropine pretreatment. **f.** The bar graph summarizes the results detailed above (numbers of measured neurons are in parenthesis). **=p<0.01, ***=p<0.005. Horizontal black lines above the recordings show the presence of various AChR receptor subtype-specific inhibitors (mecamylamine, atropine) in the aCSF. The arrow shows the time of administration of carbachol or muscarine. Zoomed 2.5-minute-long recordings of control (upper graph) and treated (lower graph) periods are besides the recordings. Changes in the frequency of mPSCs in individual GnRH neurons are shown besides these zoomed recordings. The frequency distribution of the mPSCs is graphed under each recording. carba=carbachol, atro=atropine, musca=muscarine and meca=mecamylamine.

To confirm the regulatory role of mAChRs in the decrease in the frequency of mPSCs, in the next experiment, the mAChR agonist muscarine (100 μM) was added to the aCSF in a single bolus. The measurements revealed that no inward current was evoked by muscarine. Nevertheless, a significant decrease in the frequency of mPSCs was detected (68.1 ± 6.67% of the control value, **Fig. 6d**). However, atropine pretreatment abolished this change in frequency (103.5 ± 5.11% of the control value, **Fig. 6e**). These results of the frequency changes are summarized in the bar graph (**Fig. 6f**) showing that mAChRs play a pivotal role in the decrease in frequency of mPSCs.

#### ACh and carbachol change the firing of GnRH neurons

Postsynaptic currents and firing activity of GnRH neurons positively correlate with each other (Chu and Moenter, 2005; Christian and Moenter, 2007). To evaluate the effect of acetylcholine (ACh) on the electric activity of GnRH neurons, firing was recorded in whole-cell current clamp mode. A single bolus of ACh (50 μM), the natural ligand of the cholinergic system, evoked a biphasic change in the firing rate (**Fig. 7a**). An initial stimulatory phase was followed by a robust inhibitory phase. A thorough examination of the recording, however, revealed that the stimulatory phase can be further divided into two subphases (phase I = 0.5 min, phase II = 3 min), while no subphases could be defined in the inhibitory phase (phase III = 3 min). The length of these phases was defined by an approximate estimation and did not take into consideration accidental overlaps. The firing rate changed significantly in all three phases (**Fig. 7a**, phase I: 409.3 ± 49.44%, phase II: 285.3 ± 39.91%, phase III: 44.6 ± 8.93%, washout: 98.7 ± 10.18% of the control firing rate).

**Fig. 7.**
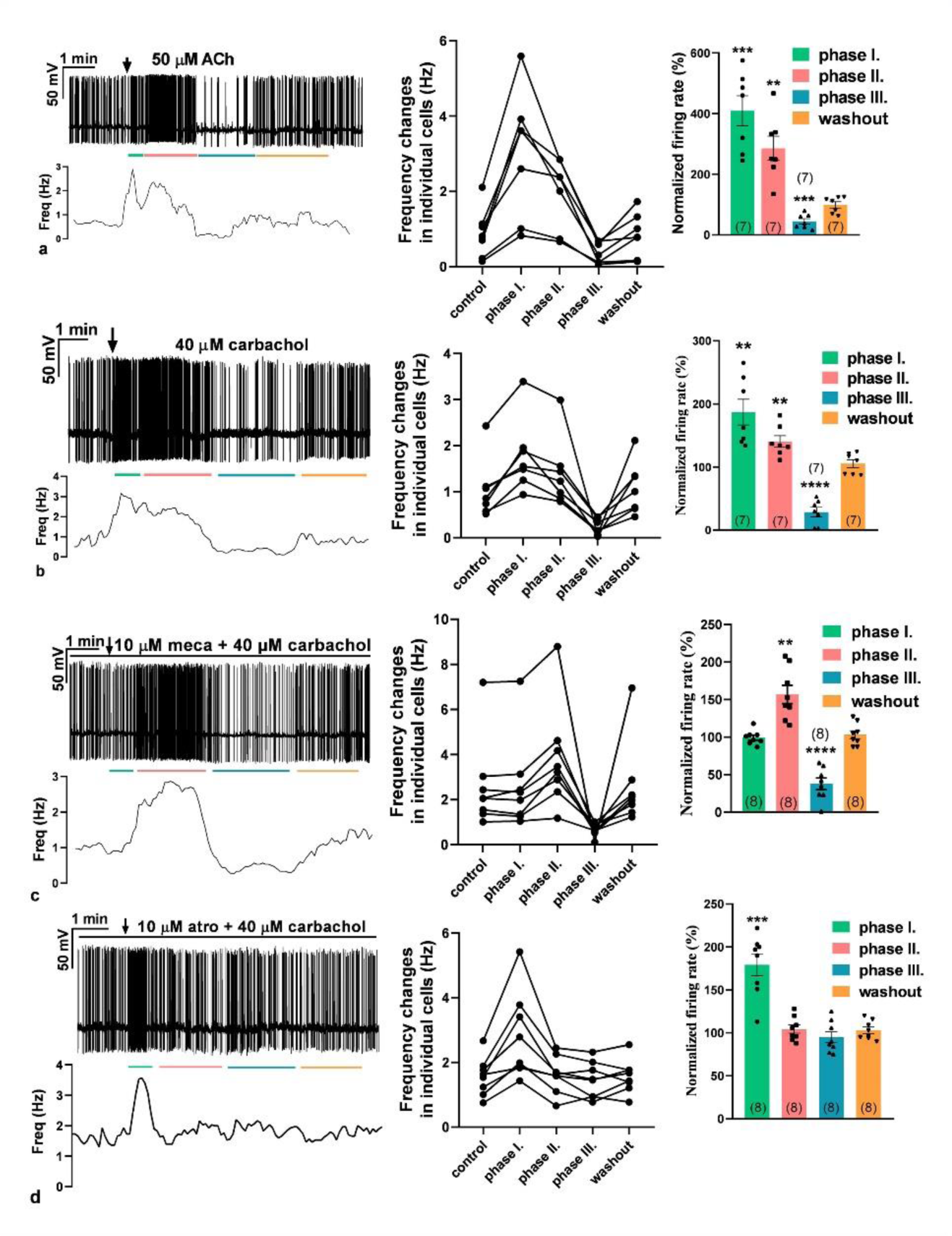
Effect of acetylcholine and carbachol on the firing rate of GnRH neurons. **a.** ACh (50 μM) evoked a biphasic action. First, it increased, then robustly inhibited the firing rate. **b**. Carbachol (40 μM) also triggered a biphasic effect with a first elevation followed by a later decline in the firing rate. **c**. Mecamylamine (10 μM, nicotinic acetylcholine receptor antagonist) pretreatment eliminated the first, 0.5-minute part of the elevation phase, suggesting that the facilitatory phase can be further divided into two subphases standing for nicotinic (phase I) and muscarinic (phase II) stimulatory actions, respectively. **d.** Atropine (10 μM) pretreatment abolished both phase II of facilitation and the inhibitory phase (phase III), proving that these phases stand for muscarinic effects. The arrow shows the time of administration of ACh or carbachol. The horizontal black line above the recordings indicates the presence of inhibitors (mecamylamine or atropine). Changes in the firing rate of individual GnRH neurons and the percentage changes are shown besides each recording. Frequency distribution of firing rate is graphed under each recording. Phase I = 0.5 min, phase II = 3 min, phase III = 3 min. The colored lines under the recordings match the colors of the bar graphs and represent the various phases. **=p<0.01, ***=p<0.005, ****=p<0.001. Meca=mecamylamine, atro=atropine. The numbers of the measured neurons are in parenthesis.

We also studied the effects of carbachol (a widely used AChR agonist not metabolized by ACh-esterase) administration (40 μM) on the firing rate of GnRH neurons (**Fig. 7b**). The phases of the evoked effects mimicked those of ACh and showed significant changes in the firing rate (**Fig. 7b**, phase I: 186.9 ± 20.47%, phase II: 140.4 ± 9.23%, phase III: 28.6 ± 7.486%, washout: 105.1 ± 6.27% of the control firing rate [the data before agonist application]).

To elucidate the nature of AChRs involved in the observed effect, the brain slices were pretreated with mecamylamine (10 μM) then carbachol was applied. Carbachol application first accelerated and then inhibited the firing activity of GnRH neurons (**Fig. 7c**). The examination of the recordings, however, clearly showed that phase I was absent, whereas phase II was present in the stimulatory phase, showing that functional nAChRs play a role in phase I. Statistical analysis revealed that phase II and phase III represented significant changes in the firing rate (**Fig. 7c**, phase I: 99.4 ± 3.311%, phase II: 156.9 ± 12.16%, phase III: 37.7 ± 7.67%, washout: 103.4 ± 5.29% of the control firing rate), suggesting that mAChRs also take part in the cholinergic control of GnRH neuron firing.

To confirm the contribution of mAChRs to the control of GnRH neuron firing, the brain slices were pretreated with atropine (10 μM), and then carbachol was added in a single bolus to the aCSF in the measuring chamber. The recordings showed that carbachol induced a short stimulatory phase, but no subsequent inhibition occurred in the presence of atropine (**Fig. 7d**). The short stimulation resembled phase I. In contrast, phases II and III were absent (**Fig. 7d**, phase I: 179.3 ± 12.6%, phase II: 104.5 ± 4.95%, phase III: 94.9 ± 6.485%, washout: 102.9 ± 3.912% of the control firing rate).

Altogether, these results show that nAChRs control phase I of the evoked firing, while phases II and III are regulated by stimulatory and inhibitory subtypes of mAChRs, respectively.

#### Electrophysiological characterization of functional nicotinic receptor subtypes involved in the control of GnRH neurons

The qRT-PCR study revealed the expression of genes encoding various subunits of nAChR in GnRH neurons. Whole-cell voltage clamp measurements were carried out to identify the functional subtypes of nAChR involved in the control of GnRH neurons. Application of nicotine (10 μM, broadband agonist of nAChRs) triggered a robust inward current (amplitude = −41.5 ± 9.05 pA, duration = 58.9 ± 18.3 s, **Fig 8a and 8d**). One minute after the inward current had decayed, nicotine was applied again. The amplitude and duration of the second response to nicotine were the same as in its first application, indicating that desensitization does not occur at this time scale (not shown). Then the slice was pretreated with DHBE (1 μM, antagonist of the α4β2 nAChR), and nicotine was applied again to the same neuron. An inward current was evoked again; however, its amplitude was significantly lower compared to the one observed when nicotine was added alone (amplitude = −18.4 ± 3.41 pA, duration = 84.4 ± 33.41 s, **Fig. 8b and 8d**), indicating that the α4β2 subtype of nAChR plays a role in the observed effect. Next, α-conotoxin AuIB (1 μM, antagonist of α3β4 nAChR) was added to the aCSF containing DHBE, and nicotine was administered again to the same neuron. This inhibitor cocktail significantly decreased the amplitude further (amplitude = −9.2 ± 2.36 pA, duration = 78.2 ± 38.6 s, **Fig. 8c and 8d**), although it did not abolish the response completely. This suggests that GnRH neurons that express the α4β2 subtype also present the α3β4 subtype of nAChR.

**Fig. 8.**
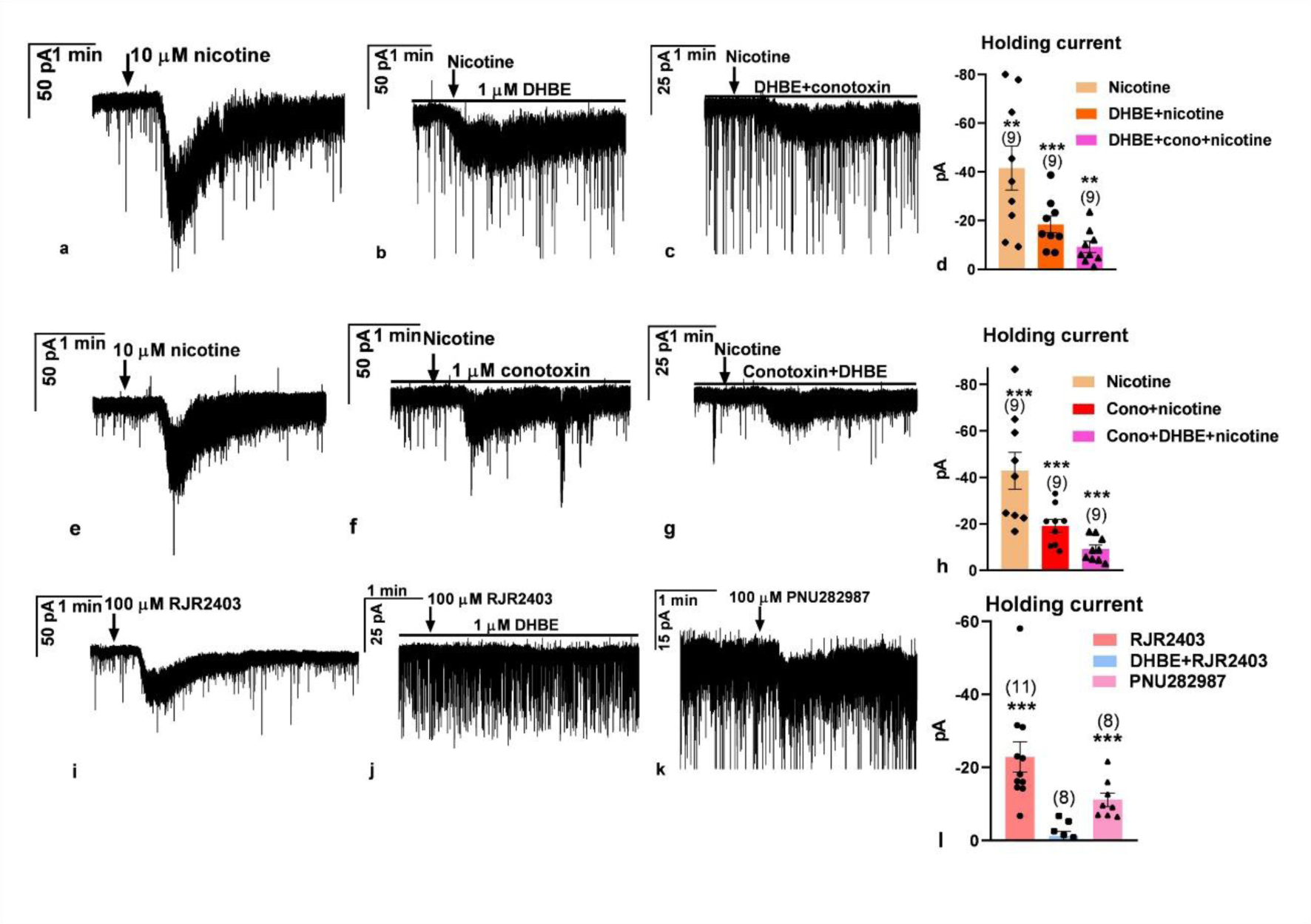
Characterization of functional nicotinic ACh receptors in GnRH neurons. Representative whole-cell patch voltage clamp recordings demonstrate that nicotine evokes an inward current in GnRH neurons, depending on the specific type of the ionotropic AChR. **a.** Nicotine administration (10 μM) induced a robust inward current. **b.** The α4β2 nAChR antagonist, DHBE (1 μM), significantly reduced the nicotine-evoked inward current measured in the same neuron as in **a** and **c**. The co-application of the α3β4 antagonist conotoxin (1 μM) with DHBE dampened the nicotine-evoked inward current further but did not eliminate it. The recording was carried out in the same neuron as in **a** and **b,** showing that neurons expressing the α4β2 receptor also present the α3β4 receptor. **d.** The bar graph summarizes the significant inward currents measured. **e.** In another neuron, nicotine also induced a strong inward current. **f.** The α3β4 receptor antagonist, conotoxin (1 μM), significantly reduced the evoked inward current measured in the same neuron as in **e** and **g**. The co-administration of DHBE attenuated the inward current further but did not abolish it. The recording was measured in the same neuron as in **e** and **f,** showing that neurons expressing the α3β4 receptors also have functional α4β2 receptors. **h**. The bar graph summarizes these results. **i-j.** The α4β2 receptor agonist, RJR2403 (100 μM), also triggered an inward current which was abolished by DHBE pretreatment. **k**. The α7 subtype nAChR agonist, PNU282987 (100 μM) also evoked a low-amplitude inward current. **l.** The bar graph summarizes these results. DHBE=dihydro-β-erythroidine (α4β2 antagonist), conotoxin=α-conotoxin AuIB (α3β4 antagonist), RJR2403 (α4β2 agonist), PNU282987 (α7 agonist). The arrow shows the time of administration of various drugs. *=p<0.05, **<0.01, ***=p<0.005. The recordings in **a-b-c** were carried out in the first slice, and the recordings in **e-g-f** were carried out in another slice.

To confirm this co-expression further, we next examined whether GnRH neurons expressing the α3β4 nAChR subtype also express the α4β2 subtype. Therefore, after measuring the nicotine-evoked inward current (amplitude = −42.9 ± 7.89 pA, duration = 68.7 ± 21.3 s, **Fig. 8e and 8h**), α-conotoxin AuIB was first added to the aCSF, and the nicotinic response was recorded again in the same neuron. Conotoxin pretreatment significantly decreased but did not eliminate the amplitude of the inward current (amplitude = −19.2 ± 2.83 pA, duration = 74.1 ± 28.39 s, **Fig. 8f and 8h**), showing expression of the functional α3β4 type receptor. Then DHBE was also added to the aCSF. This cocktail significantly lowered the amplitude further but did not abolish the evoked response (amplitude = −9.2 ± 1.74 pA, duration = 71.1 ± 26.98 s, **Fig 8g and 8h**).

To present further evidence that the α4β2 subtype of nAChR is expressed in GnRH neurons, RJR2403 (100 μM, selective agonist of the α4β2 receptor) was applied in a single bolus. Application of RJR2403 evoked an inward current (amplitude = −22.9 ± 4.15 pA, duration = 98.9 ± 38.15 s, **Fig. 8i and 8l**), which was abolished completely by DHBE pretreatment (amplitude = −1.2 ± 1.23 pA, **Fig. 8j and 8l**). Because simultaneous blockade of α4β2 and α3β4 subtypes of nAChR did not completely eliminate the nicotine-evoked response in GnRH neurons, the participation of further nicotinic receptor(s) seemed reasonable in the regulatory process. We challenged the putative expression of the α7 subtype of the receptor. Therefore, PNU282987 (100 μM, selective agonist of the α7 subunits) was applied into the bath in a single bolus. It evoked a low amplitude but still significant inward current (amplitude = −11.2 ± 1.88 pA, duration = 88.6 ± 25.15 s, **Fig 8k and 8l**). This finding indicates that the α7 subtype of nAChR also contributes to the regulation of GnRH neurons, in addition to the α4β2 and the α3β4 receptor subtypes.

### Characterization of muscarinic receptor subtypes controlling GnRH neuron activity

#### Regulation of mPSC frequencies

Using whole-cell voltage clamp recordings, we examined the contribution of stimulatory (M1, M3) and inhibitory (M2, M4) muscarinic receptors to the regulation of mPSC frequencies in GnRH neurons. We have to note that mPSCs are exclusively GABAergic (Bhattarai et al., 2011) and GABA is excitatory via GABA_A_-R due to the high intracellular chloride concentration in GnRH neurons of male mice (Herbison and Moenter, 2011). Following a 1-min-long control registration period, blockade of the inhibitory receptors was carried out with a cocktail consisting of the M2 and M4 muscarinic receptor inhibitors, tropicamide and methoctramine, respectively. The treatment significantly increased the frequency of the mPSCs, suggesting a tonic release of ACh (127.3 ± 8.50% of the control value, **Fig. 9a and 9e**). Then, muscarine (100 μM) was added in a single bolus into the measuring chamber in the continuous presence of the blocker cocktail. Muscarine significantly increased the frequency further (139.1 ± 14.75% of the pre-treatment value, **Fig 9a and 9f**). In the second step, the M1 and M3 stimulatory receptors were antagonized with a cocktail of darifenacin and pirenzepine, inhibitors of the M1 and M3 muscarinic receptors, respectively, and the measurements were carried out again. Blockade of the stimulatory muscarinic receptors decreased the frequency of the mPSCs significantly, confirming the tonic release of ACh (80.7 ± 2.67% of the control value, **Fig. 9b and 9e**). Muscarine administration significantly diminished the frequency further (139.1 ± 14.75% of the pre-treatment value, **Fig 9b and 9f**).

**Fig. 9.**
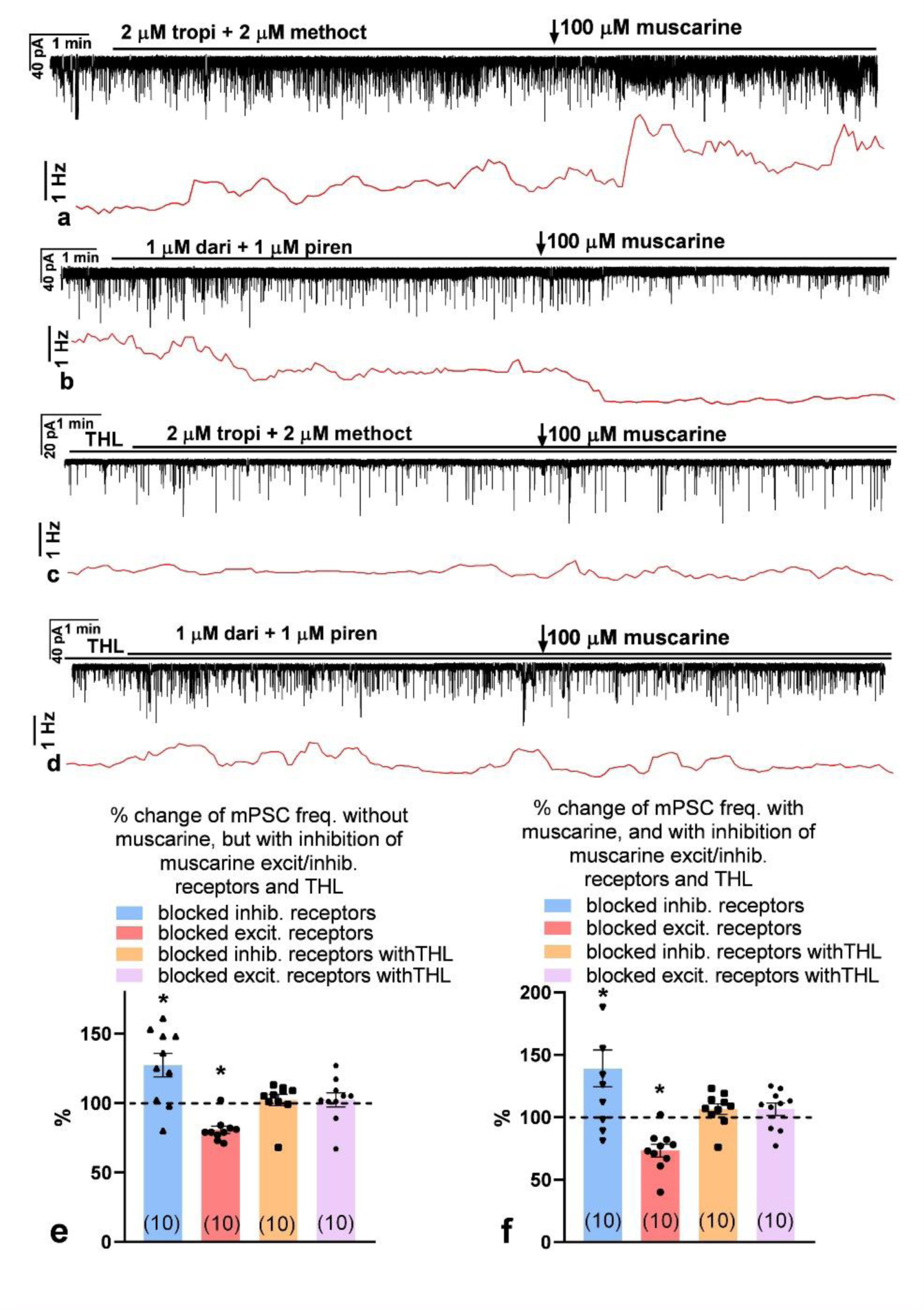
Functional characterization of muscarinic ACh receptors in GnRH neurons. Representative recordings prove the effect of blockade of inhibitory (tropicamide+methoctramine) or stimulatory (darifenacin+pirenzepine) muscarinic receptors and muscarine on mPSCs in GnRH neurons in the intracellular absence or presence of THL, an inhibitor of retrograde 2-AG endocannabinoid signaling pathway. **a.** Blockade of inhibitory receptors increased frequency of mPSCs. Application of muscarine elevated the frequency further. **b.** Conversely, blockade of stimulatory receptors diminished the frequency of mPSCs, and muscarine reduced it further. **c-d.** Intracellular presence of THL, however, abolished the effects; neither blockade of inhibitory and stimulatory receptors, nor application of muscarine evoked any significant action on the frequency. **e-f.** The bar graphs summarize these significant actions (numbers of measured neurons are in parenthesis). Horizontal black lines above the recordings show the presence of various inhibitors (THL, dari=darifenacin [M3 inhibitor], methoct=methoctramine [M2 inhibitor], piren=pirenzepine [M1 inhibitor], tropi=tropicamide [M4 inhibitor]) and arrow points the time of muscarine administration). *=p<0.05. Frequency distribution graphs are found under each recording.

Our earlier studies revealed that activation of retrograde 2-AG endocannabinoid signaling pathways can change the frequency of mPSCs in GnRH neurons. Therefore, tetrahydrolipstatin (THL), an inhibitor of 2-AG endocannabinoid production, was applied intracellularly in the patch pipette solution. Under this condition, a cocktail of tropicamide and methoctramine evoked no notable change in the frequency of the mPSCs (102.2 ± 4.1% of the control value, **Fig. 9c and 9e**), and muscarine application triggered no observable change either (106.5 ± 4.17% of the control value, **Fig 9c and 9f**). Similarly, a cocktail of darifenacin and pirenzepine resulted in no notable change in the frequency in the intracellular presence of THL (102.3 ± 5.1% of the control value, **Fig. 9d and 9e**) and muscarine induced no significant alteration of the frequency either (106.4 ± 4.97% of the control value, **Fig 9d and 9f**). These data unveil the regulatory role of retrograde 2-AG endocannabinoid signaling in the modulation of GnRH neuron physiology via both inhibitory and stimulatory types of muscarinic receptors.

#### Muscarine exerts a biphasic effect on firing of GnRH neurons

Our results suggested that besides the nACh-Rs, functional mACh-R subtypes also exist in GnRH neurons. Therefore, the putative effects of the M1-M4 receptor forms on the firing activity of GnRH neurons were studied using the whole-cell current clamp method. When muscarine (100 μM, broad-band agonist of mAChRs) was administered in a single bolus, it evoked a biphasic effect. In the first 3-minute period (phase I), muscarine significantly elevated the firing rate (147.6 ± 10.28% of the control firing rate, **Fig. 10a**). In the second 3-minute period (phase II), a significant decrease was seen in the firing rate (54.5 ± 6.9% of the control firing rate, **Fig. 10a**). To further dissect the phenomenon, first, we examined the function of M4 by pretreating the brain slices with 2 μM tropicamide, a selective inhibitor of the M4 subtype of the mAChR, then muscarine was administered. The recording clearly showed that both the stimulatory and inhibitory phases developed (**Fig. 10b**). The firing rate significantly increased up to 152 ± 14.8% of the control value and then significantly decreased to 78.3 ± 5.53% of it. However, phase II represented a decrease that was milder than without tropicamide, suggesting that, beside M4, another inhibitory subtype is involved. Next, M2 was blocked by adding 2 μM methoctramine, a selective inhibitor of the M2 subtype of mAChR, then muscarine was applied. The stimulatory phase I persisted in the presence of methoctramine, revealing a significant increase in the firing rate (145.6 ± 6.59% of the control value, **Fig.10c**). The inhibitory phase II was also present because the firing rate significantly diminished (80.8 ± 3.45% of the control value, **Fig. 10c**). Phase II, however, displayed a smaller decrease than in the absence of the M2 or M4 antagonists, revealing the role of the M2 receptor subtype.

**Fig. 10.**
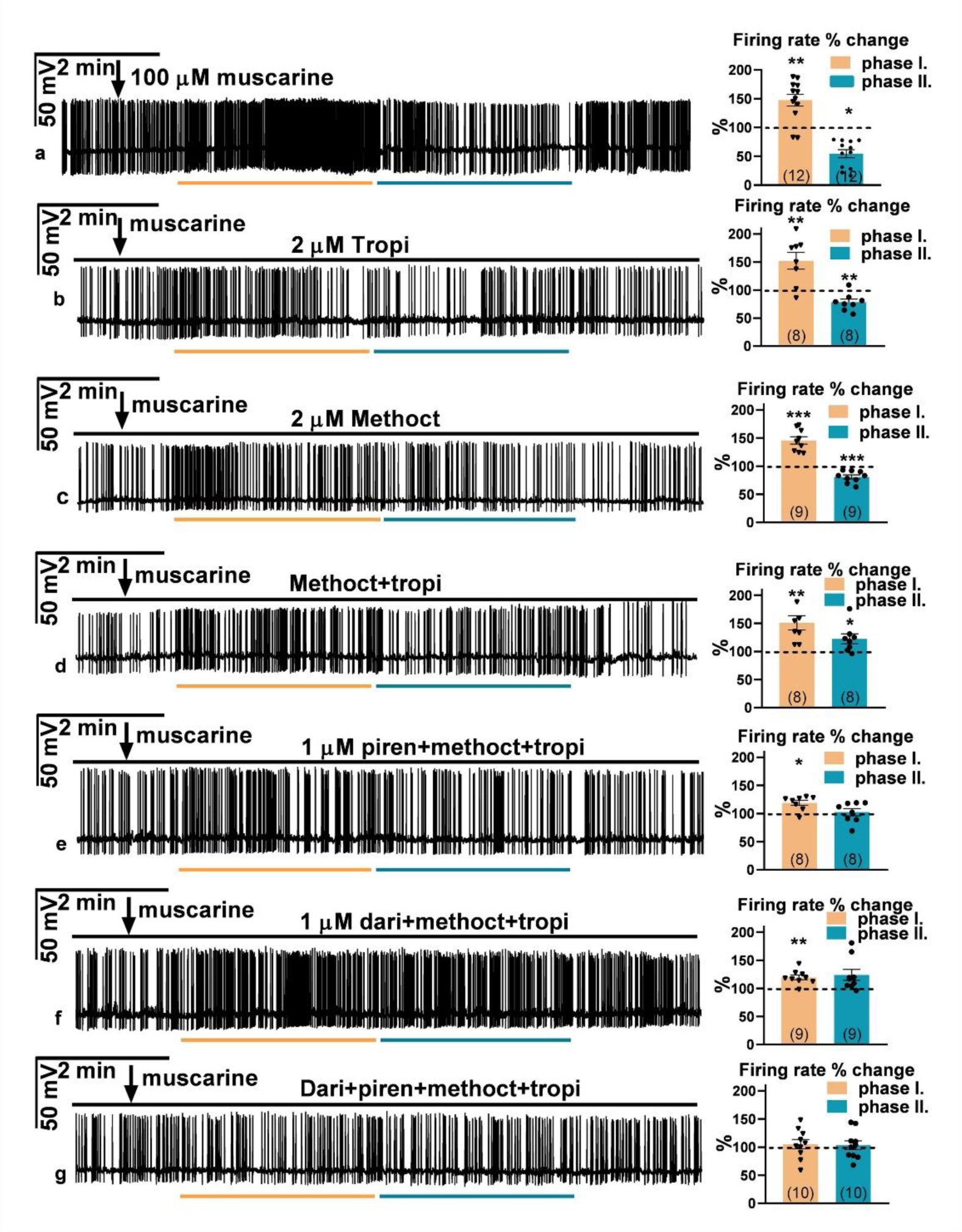
Impact of muscarine receptor subtypes on firing of GnRH neurons. Representative whole-cell patch current clamp recordings show that muscarine evoked a biphasic effect on the firing of the GnRH neurons expressing various muscarine receptor subtypes. **a**. In the first, ∼2.5-minute period, muscarine (100 μM) enhanced (phase I) the firing rate of the GnRH neuron followed by a decreased activity period (phase II). **b.** The inhibitory M4-muscarine receptor antagonist tropicamide (2 μM) dampened the inhibitory effect of muscarine (as seen in phase II). **c.** The inhibitory M2-muscarine receptor antagonist methoctramine (2 μM) also decreased the inhibitory effect of muscarine (phase II). **d**. Co-administration of tropicamide and methoctramine eliminated the inhibitory effect of muscarine (phase II). **e**. The stimulatory M1-muscarine receptor antagonist, pirenzepine (1 μM), dampened the stimulatory effect of muscarine (as seen in phase I) in the presence of the M2 and M4 antagonists. **f.** The stimulatory M3-muscarine receptor antagonist, darifenacin (1 μM), also attenuated the stimulatory effect of muscarine (phase I). **g**. Neither facilitation nor inhibition was seen when a cocktail of all four antagonists was applied. The black line above each recording shows the presence of cocktails of various muscarine receptor subtype inhibitors in the aCSF (dari=darifenacin, methoct=methoctramine, piren=pirenzepine, tropi=tropicamide). The light brown line under each recording presents period the of stimulatory phase (phase I), and blue line stands for that of the inhibitory phase (phase II). Arrows show the administration of muscarine. *=p<0.05, **<0.01, ***=p<0.005.

Next, we antagonized both inhibitory subtypes. In the presence of a cocktail of tropicamide and methoctramine, the stimulatory phase I still existed (150.8 ± 12.61% of the control firing rate), but no inhibitory phase II developed; rather, stimulation was seen in phase II period (122.4 ± 8.76% of the control value) (**Fig. 10d**). The changes in both phases were significant. The stimulatory period, therefore, lasted much longer in the presence of the M2 and M4 antagonists.

We also examined the function of the stimulatory mACh-R subtypes, M1 and M3. First, pirenzepine (1 μM, selective antagonist of the M1 subtype of mAChR) was added to the cocktail of tropicamide and methoctramine, and then muscarine was applied. In the presence of these three antagonists, muscarine significantly increased the firing rate in phase I (118.9 ± 4.5% of the control firing rate, **Fig. 10e**). Nevertheless, the value of the increase was lower than without pirenzepine, showing the role of M1 in the stimulation. Phase II, however, presented no significant change at all (102.4 ± 6.42% of the control value). Thus, the duration of the stimulation was not longer than three minutes.

Role of M3, the other stimulatory mAChR subtype, was also investigated. Darifenacin (1 μM, selective antagonist of the M3 subtype of mAChRs) was added to the cocktail of tropicamide and methoctramine, and then muscarine was applied. Phase I. presented a significant elevation in the firing rate (119.4 ± 4.18% of the control firing rate, **Fig. 10f**. Phase II. also displayed a significant increase in the firing rate (124.0 ± 9.71% of the control value, **Fig. 10f**, thus the stimulation lasted much longer than three minutes.

Finally, the effect of muscarine was examined in the presence of M1-M2-M3-M4 antagonists. This cocktail abolished both phase I. (105.3 ± 8.31% of the control value) and phase II. (103.2 ± 7.89% of the control firing rate) completely (**Fig. 10g**).

#### Optogenetic activation of cholinergic afferents changes the electrophysiological activity of GnRH neurons and reveals co-transmission of ACh and GABA

To study the effect of endogenous ACh, GnRH neurons of triple-transgenic Chat-Cre-ChR2-GnRH-GFP mice were examined in acute brain slices using LED square pulses (470 nm, 5 ms) to activate channelrhodopsin-expressing cholinergic axons. GnRH neurons were whole-cell clamped in current-clamp mode at 0 pA to detect firing. After a 30-second control period, an LED train of 5 Hz was applied for a subsequent 60 second while firing was continuously recorded. A biphasic pattern was detected when a 5 Hz train was applied: the initial stimulation (191.2 ± 33.19%, p=0.0157, t=2.748, df=14, t-test) was followed by a transient inhibition (71.0 ± 9.02%, p=0.0063, t=3.212, df=14). Stimulation lasted for 15 ± 2.7 s; inhibition lasted for 38 ± 3.4 s (**Fig. 11a and 11c**.). Both stimulation and inhibition disappeared in the presence of a cocktail of atropine and mecamylamine (**Fig 11b and 11c**.), suggesting the involvement of ACh in the process.

**Fig. 11.**
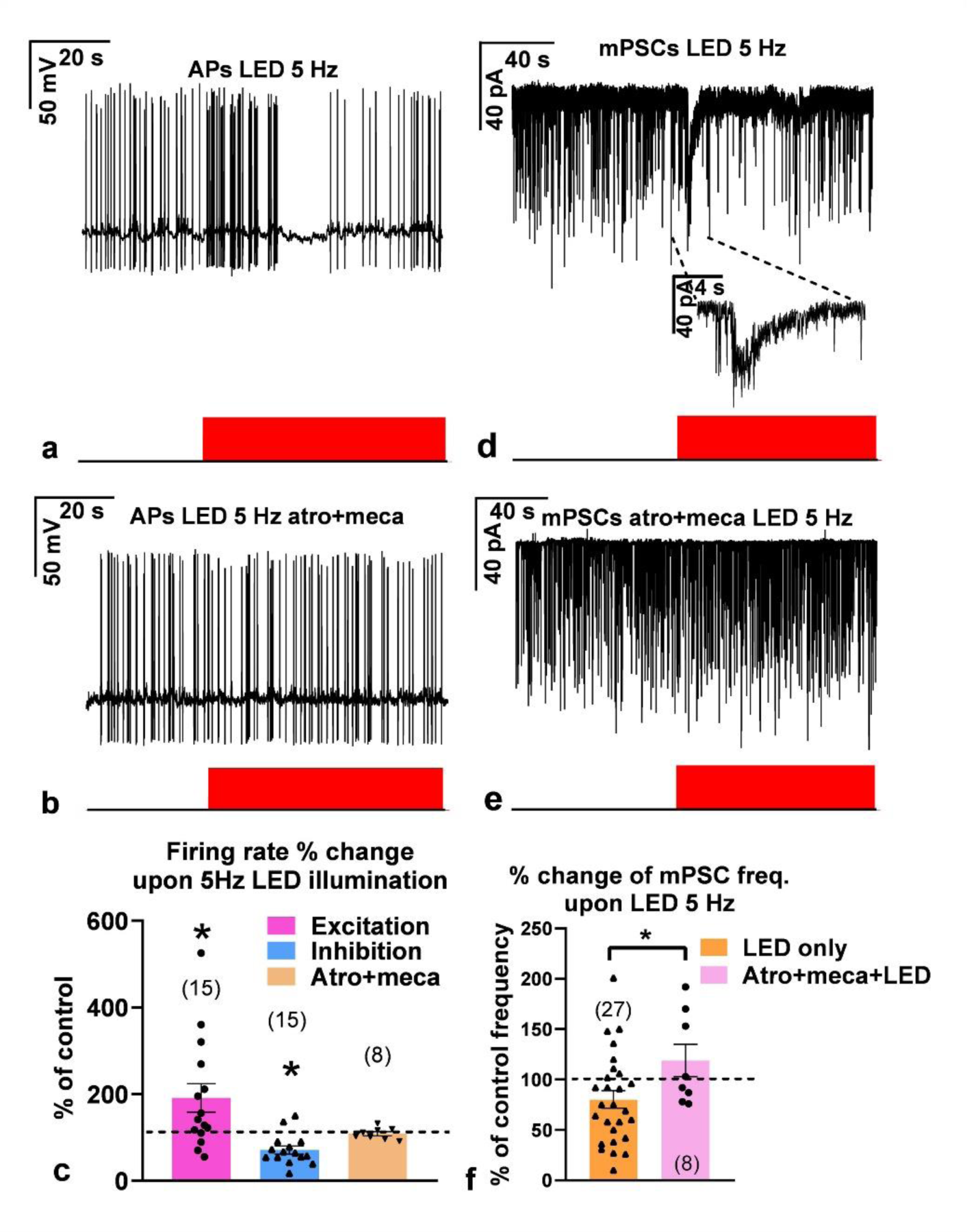
Effects of optogenetic stimulation of cholinergic axons upon firing activity and frequency of spontaneous mPSCs of GnRH neurons. **a.** LED illumination at 5 Hz evoked an initial facilitation followed by transient inhibition in firing. **b**. Both the stimulatory and inhibitory effects were eliminated when the slices were pretreated with the cocktail of atropine and mecamylamine. **c**. The bar graph summarizes the effect of LED illumination at 5 Hz, revealing that ACh was released from channelrhodopsin-enriched cholinergic terminals. **d.** LED illumination at 5 Hz triggered an inward current, followed by a transient decrease in the mPSC frequency. The inset is a zoomed-in region of the graph containing the inward current. **e.** Both the inward current and the diminished mPSC frequency disappeared in the presence of a cocktail of atropine and mecamylamine. **f.** The bar graph shows a significant difference in the mPSC frequencies measured in the absence or presence of the inhibitor cocktail. Atro=atropine, meca=mecamylamine. Recordings are in black; red bars underneath the recordings indicate the period of LED illumination. The numbers of the measured neurons are in parentheses. *=p<0.05.

The effects of optogenetic activation of cholinergic afferents were seen in the membrane currents, too. After a 2-minute control period, LED illumination at 5 Hz for 2 minutes induced an initial inward current (amplitude: 51 ± 12.7 pA; **Fig. 11d and 11f**.) resembling those triggered by carbachol. A transient decrease in the frequency was then observed (80.2 ± 8.65% of the control value, n = 27; **Fig. 11d and 11f**). Both the inward current and the decreased frequency were eliminated in the presence of atropine+mecamylamine (frequency: 118.9 ± 16.16% of the control, n=8; **Fig. 11e and 11f**). The difference in the frequency data was significant (Student’s t-test, p=0.0405, df=33, t=2.132).

To examine the extra- and/or synaptic character of cholinergic neurotransmission onto GnRH neurons, the existence or absence of evoked mPSC (emPSC) was investigated in the presence of TTX in GnRH neurons of the adult male Chat-Cre-ChR2-GnRH-GFP transgenic mice. LED illumination (0.2 Hz, 470 nm, 5 ms duration, then averaged of 60 runs of the 5 s long recordings) triggered no visible emPSC response in 90 % of GnRH neurons suggesting an extra-synaptic action of ACh. In 10% of the cells, however, robust emPSC was seen (**Fig 12a**). The emPSCs occurred synchronously with the LED square pulse (**Fig 12, zoomed inset**). The onset was 2-3 ms after the rise of the illumination. The emPSC was still observable in the same GnRH neuron in the presence of atropine and mecamylamine (10-10 µM, **Fig. 12b**) showing the release of a neurotransmitter or neuromodulator other than ACh.

**Fig. 12.**
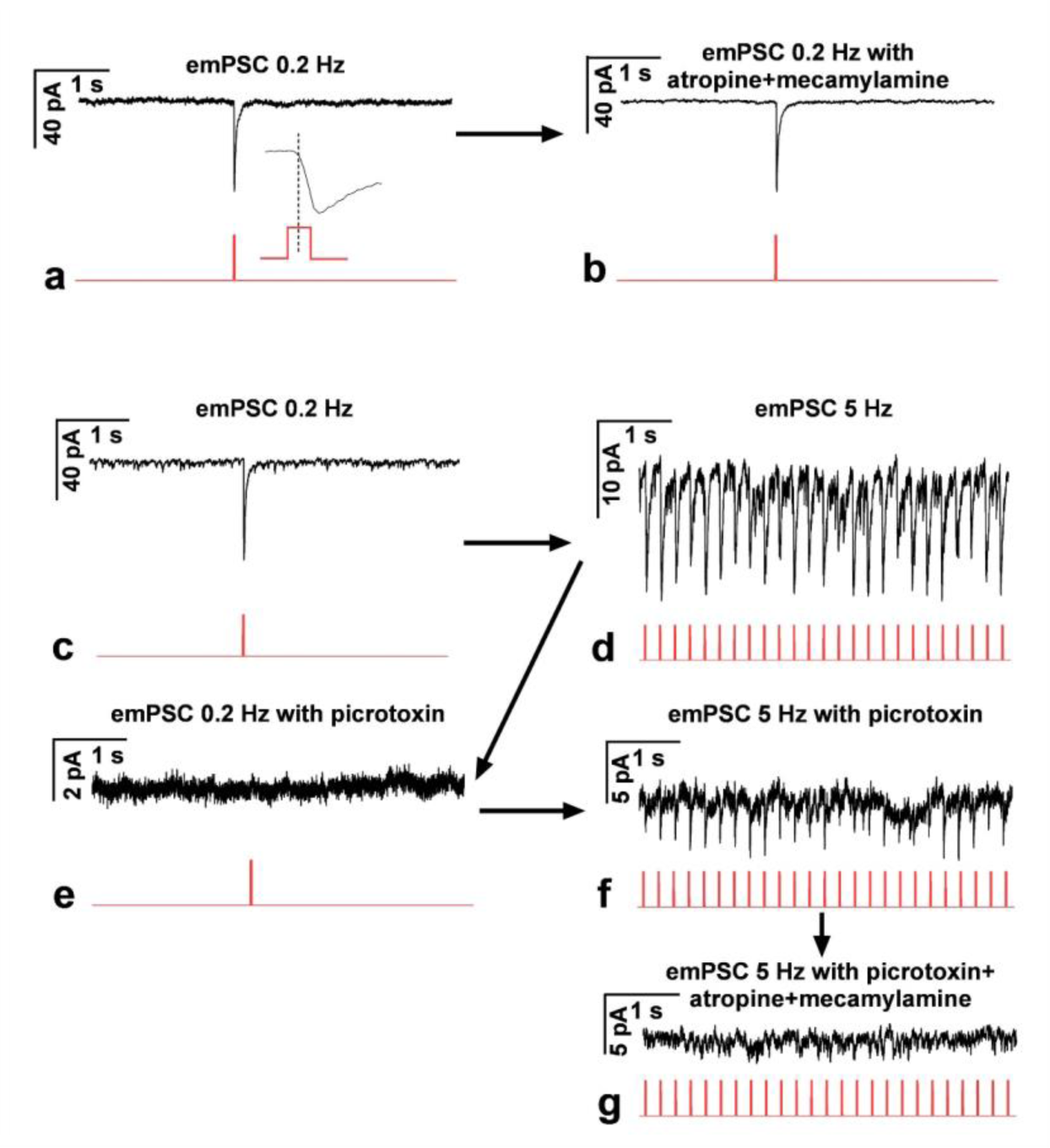
Evoked mPSCs (emPSCs) observed in a subpopulation of GnRH neurons of Chat-Cre-ChR2-GnRH-GFP transgenic mice upon LED illumination at 0.2 or 5 Hz. **a.** A robust inward emPSC was triggered by LED illumination at 0.2 Hz. **b**. The emPSC was still seen in the presence of atropine and mecamylamine in the same neuron as in **a**. **c**. A similar emPSC was revealed in another GnRH neuron at 0.2 Hz. **d.** A train of emPSCs of lower amplitude was evoked by LED at 5 Hz in the same neuron as in **c**. **e.** The emPSC observed at 0.2 Hz was eliminated by picrotoxin in the same neuron as in **d**. **f.** A train of emPSCs was still detected in the presence of picrotoxin at 5 Hz in the same neuron as in **e**. **g**. This train was abolished by adding atropine and mecamylamine to the picrotoxin-containing aCSF in the same neuron as in **f**. Arrows show the time order of measurements in the same neuron. Recordings are in black; red lines underneath the recordings indicate the period of LED illumination at 0.2 or 5 Hz, respectively.

In other GnRH neurons of the triple transgenic mouse in which emPSC was evoked by LED of 0.2 Hz (**Fig. 12c**), we examined the effect of a 5 Hz LED illumination (5 Hz train, then averaged of 60 runs of the 5 s long recordings). In 50% of these neurons, a train of emPSCs was seen (**Fig. 12d**). These emPSCs were synchronous with the 5 Hz LED pulses, although the amplitude was lower than the one evoked by the 0.2 Hz (≈20%). To identify the transmitter(s) responsible for the emPSCs, first picrotoxin (100 µM) was added to the aCSF, and the existence or absence of synaptic response was examined at 0.2 or 5 Hz in the same GnRH neuron as in **Fig. 12c and 12d**. Picrotoxin abolished the emPSC at 0.2 Hz completely, providing strong evidence for the exclusive release of GABA from ChR-2-tagged cholinergic axons (**Fig. 12e**) at this LED frequency. Surprisingly, a train of emPSCs was still detected at 5 Hz illumination (**Fig. 12f**) in the presence of picrotoxin in this same neuron. The amplitudes of these emPSCs were lower than those detected without the GABA-A-R blocker. The emPSCs triggered by LED at 5 Hz were eliminated when atropine-mecamylamine was added to the aCSF beside picrotoxin (**Fig. 12g**). The last two observations suggest the release of both ACh and GABA at 5 Hz. These results show that cholinergic axons innervating GnRH neurons use ACh and GABA for neurotransmission in a frequency-dependent manner.

The above optogenetic studies showed that GABA and acetylcholine are co-transmitted from a subpopulation of cholinergic axon terminals, depending on the frequency of LED stimulation. To provide the structural correlate of ACh/GABA co-transmission, immunofluorescence triple labeling was performed on preoptic sections of Chat-Cre-ChR2 transgenic mouse brains for vesicular GABA transporter (VGAT), green fluorescent protein (GFP)-tagged channelrhodopsin-2 (ChR2-GFP), and GnRH. The confocal microscopic analysis revealed that VGAT-IR (**Fig. 13a and 13b**), and cholinergic ChR2-GFP-IR (**Fig. 13a and 13c**) axons exist in the proximity of GnRH neurons. At high power, the VGAT (**Fig. 13b**) and ChR2-GFP (**Fig. 13c**) were co-localized (**Fig. 13d**) in the same cholinergic axon varicosities. Furthermore, VGAT-(**Fig 13f**) and ChR2-GFP-(**Fig 13g**) expressing double-labeled axon boutons (**Fig 13e and 13i**) were found in juxtaposition (**Fig 13e and 13i**) with GnRH dendrite (**Fig 13h and 13i**). These morphological results unveiled the dual ACh/GABA chemotype of at least a subset of cholinergic axons innervating GnRH neurons and support the optogenetic findings on ACh/ GABA co-transmission in regulation of GnRH neurons.

**Fig. 13.**
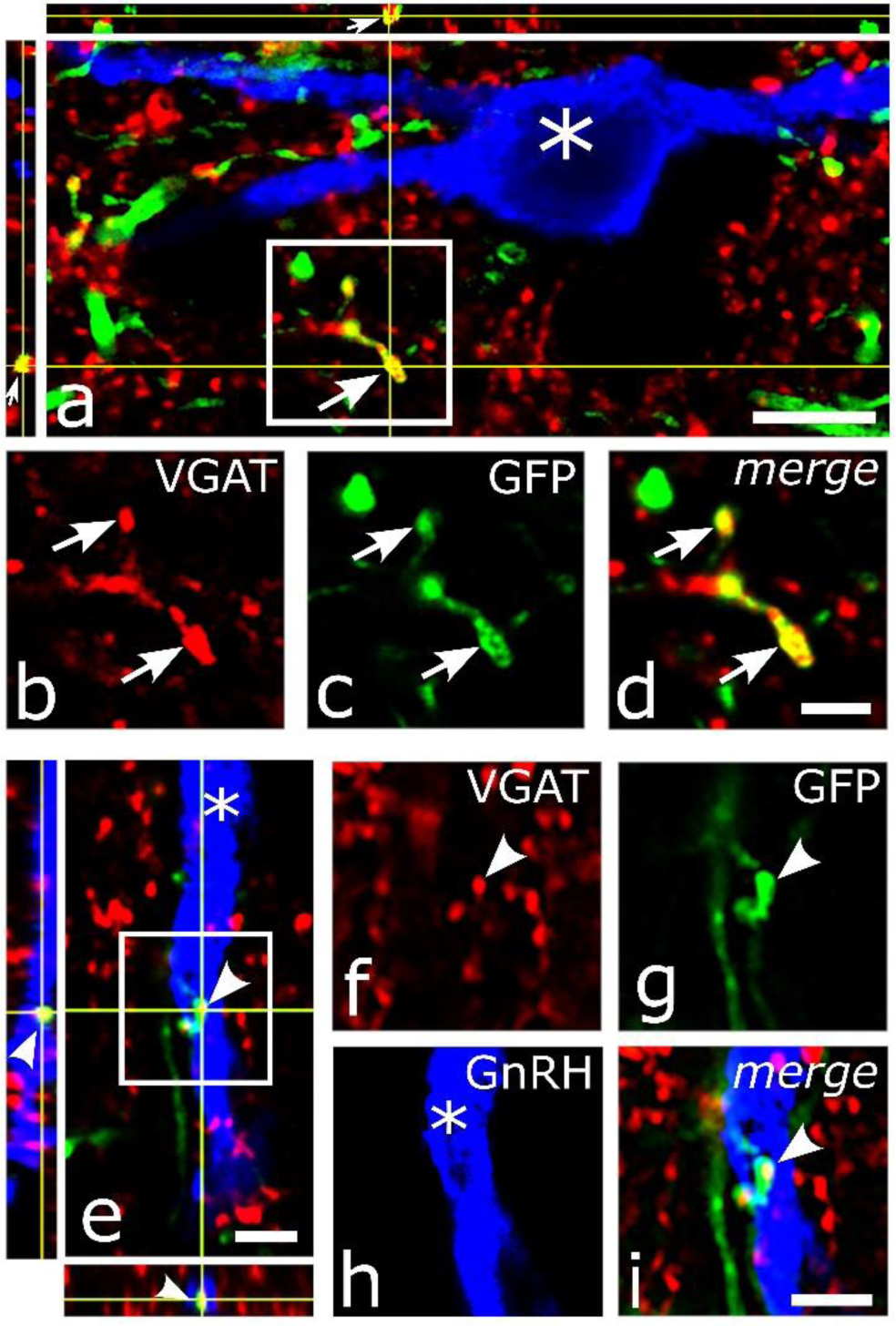
Expression of vesicular GABA transporter in cholinergic axons targeting GnRH neurons in the Chat-Cre-ChR2 mouse. **a.** Triple immunofluorescent imaging revealed vesicular GABA transporter (VGAT, red channel), GFP-tagged channelrhodopsin-2 (ChR2-GFP, green), and GnRH (blue)-immunoreactive (IR) structures via confocal microscopy. VGAT (**b**) and ChR2-GFP (**c**) were co-localized (**d**) within the same axon varicosity (enframed in **a**, enlarged in **d**, arrows) near a GnRH neuron (blue, asterisk). The juxtaposition of a VGAT (**f**, arrowhead) and ChR2-GFP (**g**) IR double-labeled axon varicosity (**i**) with the dendrite of a GnRH neuron (**h**, asterisk) was confirmed in orthogonal views (**e**; enframed area enlargement in **i**). Scale bars: a, 5 µm; b-i, 2 µm.

## Discussion

This study provides evidence for a significant role of the central cholinergic system in the regulation of GnRH neurons and LH release in adult male mice. *1*. *In vivo* pharmacogenetic stimulation of the cholinergic system caused a rapid release of LH. *2*. The cholinergic input originated from the MS and the DBB. *3*. ACh regulated GnRH neurons via a broad set of nicotinic and muscarinic receptors. *4*. The initial excitation, caused by ACh, was followed by an inhibitory phase. *5*. Optogenetic activation of cholinergic axons resulted in a frequency-dependent co-transmission of ACh and GABA in a sub-population of axons.

We used pharmacogenetics for stimulation of the cholinergic system. Activation of DREADD-expressing cholinergic neurons by CNO administration rapidly increased both basal and mean LH levels, with peaks at 30 minutes, followed by a prolonged return to the basal level. Cholinergic drugs targeting the hypothalamus have been shown to increase the secretion of LH *in vitro* (Fiorindo and Martini, 1975). Furthermore, atropine administration into the third cerebral ventricle blocked the proestrus surge of LH and FSH in rats (Libertun and McCann, 1973).

The GnRH neuronal network (Campbell et al., 2022) is surrounded by clusters of acetylcholine-synthesizing neurons in the MS and the DBB (Wu et al., 2014; Li et al., 2018), where both cholinergic and GnRH neurons often appear intermingled. The MS-DBB-mPOA region is rich in VAChT-immunoreactive axons (Schäfer et al., 1994), and our 3DISCO-based quantitative analysis unveiled that GnRH neurons are innervation targets of these cholinergic axons. Microscopic evaluation confirmed juxtapositions of VAChT-IR axons to the somatic and dendritic domains of GnRH neurons. The ratio of cholinergic axons juxtaposed to GnRH dendrites vs somata was ∼6.4:1 emphasizing the exquisite role of GnRH dendrites in communication with neurotransmitter systems (Norberg et al., 2013). In this study, 37.4 ± 5.1 percent of GnRH-IR perikarya received cholinergic inputs, indicating cholinergic control at the level of cell bodies, too. Both single and multiple cholinergic appositions were found on the surface of GnRH somata. The double immunoelectron microscopic analysis confirmed that the juxtaposed VAChT and GnRH-IR profiles formed direct contacts without interposing astrocytic processes. The cholinergic axon terminals filled with small, electron-lucent vesicles occasionally formed synapses with GnRH neurons in the MS-DBB-mPOA region, indicating sites of synaptic- and/or volume transmission (Sarter et al., 2009). A similar network has previously been reported in rats (Turi et al., 2008).

Cholinergic neurons distributed in the mouse brain (Li et al., 2018) project to the MS-DBB-MPOA. Retrograde viral labeling of cholinergic neurons wired to GnRH neurons revealed that the cholinergic afferents of the GnRH neurons arise exclusively from the MS and DBB. MS has also been claimed recently as the main source of cholinergic neuronal inputs to GnRH neurons (Shostak et al., 2023).

Bath application of ACh (50 μM) or the acetylcholinesterase-resistant AChR agonist carbachol (40 μM) evoked a biphasic effect on GnRH neurons, first elevating then decreasing the firing rate. Since the electric activity of GnRH neurons and GnRH release correlate (Ordög et al., 1997; Zhang et al., 2007; Pielecka-Fortuna et al., 2008), the result suggests that cholinergic drugs likely induce GnRH release. Similarly, ACh first activated embryonic GnRH neurons and increased their intracellular Ca^2+^ concentration, followed by subsequent 5-minute-long decline (Shostak et al., 2023). In our hands, however, the first phase of the recordings was characterized by a fast transient increase followed by a slow, prolonged rise in the firing rate. Pretreatment of slices with nAChR antagonist mecamylamine eliminated the fast phase, leaving the slow-type facilitation and the inhibitory phase untouched, suggesting the role of nAChRs in the fast action. Indeed, nicotine can induce fast depolarization and enhanced firing rate in hypocretin cells (Huang et al., 2011) and evoke GnRH release from hypothalamic fragments (Richardson et al., 1982).

Mecamylamine did not affect the slow facilitation and the inhibition of firing, suggesting that they were due to the activation of mAChRs. Indeed, application of atropine abolished both slow components while the faster, transient facilitation remained intact. Hypothalamic CRF (Mukai et al., 2020) and POMC cells (Jeong et al., 2017), goldfish (Kawai et al., 2013) and embryonic mouse GnRH neurons (Shostak et al., 2023) also respond to muscarine, supporting the current finding.

In our study, carbachol triggered an inward current, followed by a decrease in the frequency of mPSCs. Inward current indicates membrane depolarization, resulting in fast facilitation seen in firing rates. The inward current was abolished by mecamylamine, suggesting the inevitable role of nAChRs in the process. In the hypothalamus, similar nAChR-dependent activation was reported in POMC neurons (Huang et al., 2011). Embryonic mouse GnRH neurons also responded to ACh administration with a nAChR-dependent fast and transient stimulatory change in membrane potential (Shostak et al., 2023), resembling the inward current we observed. The carbachol-triggered decrease in the frequency of mPSCs after the inward current was blocked by atropine indicates the involvement of mAChRs in this process. Previous reports have also shown mAChR-dependent ion currents in goldfish (Kawai et al., 2013) and embryonic mouse GnRH neurons (Shostak et al., 2023).

Our work also revealed the subunit compositions of nAChRs; the presence of α3β4, α4β2 and α7 receptors was confirmed in GnRH neurons, of which the α3β4 and the α4β2 were localized in the same GnRH neuron. The hypothalamic presence of functional α4β2 and α7 receptors has previously been demonstrated with the major role of α7 (Huang et al., 2011) and the expression of α3 subunit shown in embryonic GnRH neurons (Shostak et al., 2023). Interestingly, the α4, α7, β2 and β4 subunits were not found in these embryonic neurons, indicating different nAChR signaling in adult *versus* embryonic GnRH neurons. Our inward ion current measurements reflect the abundance of α4β2 and α3β4 subunits in the adult, with a relatively minor role of α7.

Embryonic GnRH neurons process the ACh signal via alpha 3 and M2 and M4 receptors (Shostak et al., 2023), in contrast with the much wider set of AChRs we found in adults. The differences in signaling mechanisms of ACh in embryonic *versus* mature GnRH neurons suggest a significant reshape in the ACh receptor profile of the GnRH neurons during their postnatal differentiation.

Nicotine triggered a robust inward ion current and excitation in GnRH neurons. The responses evoked by repeated application (90 - 120 s) of nicotine presented no desensitization at this period (Huang et al., 2011; Buls Wollman and Fregosi, 2021).

We demonstrated an inverse effect of muscarine on the mPSC frequency in the presence of either M2/M4 or M1/M3 receptor antagonists, which can be explained by the involvement of different intracellular signaling pathways. M1/M3 are G_q_-coupled receptors mediating neuronal excitation, whereas the G_i/o_-coupled M2/M4 activity leads to inhibition (Brown, 2018). The M2 or M4 antagonists prolonged the stimulatory phase, indicating that M2/M4 receptors can attenuate the cholinergic excitation. Both facilitatory and inhibitory actions were abolished by the intracellular application of tetrahydrolipstatin (THL) which bloked the endocannabinoid 2-arachidonoylglycerol (2-AG) synthesis and the subsequent release of 2-AG from GnRH neurons. Involvement of retrograde 2-AG signaling in the regulation of GnRH neurons was reported earlier (Farkas et al., 2010; Bálint et al., 2016; Farkas et al., 2016; Balint et al., 2020). Furthermore, the role of endocannabinoids in mAChRs-mediated signal transduction was shown in the brain (Fukudome et al., 2004; Uchigashima et al., 2007; Martin et al., 2015; Joo et al., 2019; Tryon et al., 2023).

LED illumination of GnRH neurons at 5 Hz in Chat-Cre-ChR2-GnRH-GFP transgenic mice evoked a biphasic change in the firing rate, which was abolished by the combined administration of mecamylamine and atropine. Antagonization of nicotinic and muscarinic AChRs also prevented the inward current and the mPSC frequency decrease evoked by the 5 Hz stimuli. The results confirmed that endogenously released ACh acting on nicotinic and muscarinic AChRs is involved in these changes.

Most of the septo-hippocampal cholinergic neurons synthesize and co-transmit GABA (Takács et al., 2018). GABA is excitatory to GnRH neurons via GABA-A receptors due to the high intracellular Cl^-^ milieu (DeFazio et al., 2002; Moenter and DeFazio, 2005; Farkas et al., 2010), although the sites of its production (Moore et al., 2015; Piet et al., 2018; Silva et al., 2019) have only partially been elucidated. Using optogenetics, LED illumination of channelrehodopsin-2-tagged cholinergic afferents of GnRH neurons at 0.2 or 5 Hz resulted in a frequency-dependent evoked response. Exclusive GABA release was evoked from cholinergic axons at 0.2 Hz. In contrast, both GABA and ACh were released from axons innervating GnRH neurons at 5 Hz. ACh and GABA co-transmission was demonstrated in the hippocampus (Takács et al., 2018). Kisspeptin and GABA co-release was shown in AVPV neurons innervating GnRH neurons (Piet et al., 2018). Frequency-dependent release was also identified in hypothalamic kisspeptin neurons of mice (Liu et al., 2011; Qiu et al., 2016).

In conclusion (**Fig. 14**), the findings collectively indicate that ACh released from cholinergic afferents, originating from the MS-DBB region, excites GnRH neurons via specific nicotinic and muscarinic receptors. The activation evokes an augmented GnRH release that, in turn, increases the LH output from the anterior pituitary.

**Fig. 14.**
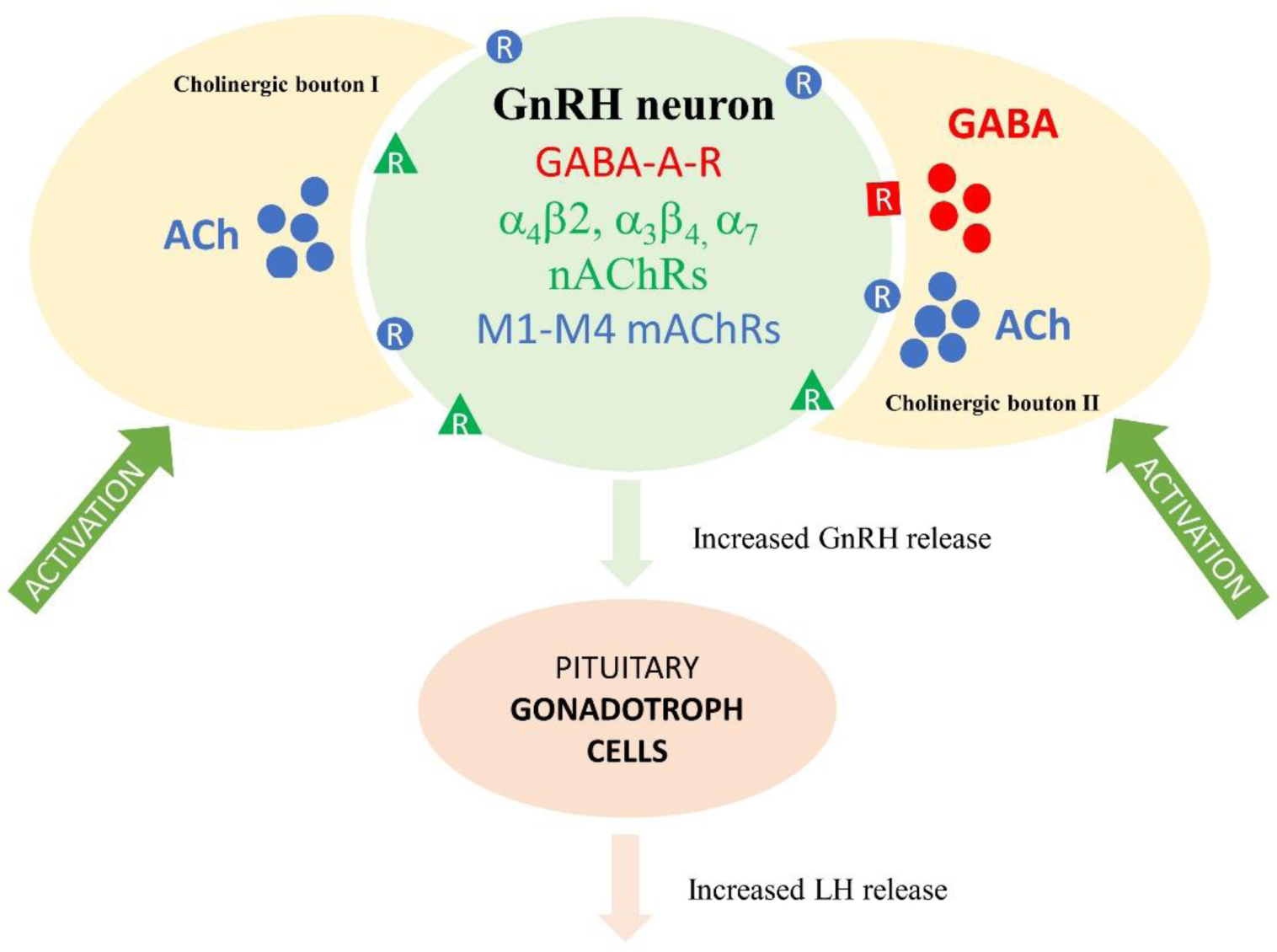
Schematic illustration of cholinergic control of GnRH neurons and the HPG axis. Cholinergic boutons (I) innervate hypophysiotropic GnRH neurons that express GABA-A receptor (GABA-A-R, red square), three subtypes (α4β2, α3β4, and α7, green triangle) of nicotinic acetylcholine receptor (nAChR), and muscarinic receptors (M1-M4, mAChR, blue oval). The released ACh utilizes both muscarinic and nicotinic receptors in communication with GnRH neurons. At least a subpopulation of cholinergic boutons (II) is capable of transmitting gamma-aminobutyric acid (GABA) in addition to acetylcholine (ACh). Activation of cholinergic boutons increases the firing of GnRH neurons, leading to increased GnRH and concurrent luteinizing hormone (LH) release.

## Acknowledgements

The authors express their gratitude to Dr. Suzanne M. Moenter (Department of Molecular and Integrative Physiology, University of Michigan, Ann Arbor, MI, USA) for the kind donation of the GnRH-GFP transgenic mice used in this study. We are also grateful to Dr. Cathrine Dulac (The Biological Labs, Harvard University, Cambridge, MA, USA) for the gift of the GnRH-Cre mouse line and to Dr. Erik Hrabovszky (Institute of Experimental Medicine, Budapest, Hungary) for the critical reading of the manuscript and the donation of the GnRH antibody. We appreciate the excellent technical assistance of Ms. Barbara Göblyös. Project no. RRF-2.3.1-21-2022-00011, titled National Laboratory of Translational Neuroscience, has been implemented with the support provided by the Recovery and Resilience Facility of the European Union within the framework of Programme Széchenyi Plan Plus.

## Abbreviations

AAV: adeno-associated virus
ACh: acetylcholine
Chat: choline acetyltransferase
ChR2: channelrhodopsin 2
CNO: clozapine N-oxide
Cono: α−conotoxin
AuIB Dari: darifenacin
DBB: diagonal band of Broca
DHBE: dihydro-β-erythroidine hydrobromide
DREADD: Designer Receptors Exclusively Activated by Designer Drug
ELISA: enzyme-linked immunosorbent assay
emPSC: evoked miniature postsynaptic current
G: G-protein
Gq: Gq protein alpha subunit
GFP: green fluorescent protein
GnRH: gonadotropin-releasing hormone
HPG: hypothalamo-pituitary-gonadal
ICC: immunocytochemistry
LH: luteinizing hormone
methoct: methoctramine
M: muscarinic receptor
mPOA: medial preoptic area
MS: medial septum
nAChR: nicotinic acetylcholine receptor
mAChR: muscarinic acetylcholine receptor
mPSC: miniature postsynaptic current
piren: pirenzepine
RJR: RJR2403
PNU: PNU282987
RV: rabies virus
THL: tetrahydrolipstatin
tropi: tropicamide
TVA: avian tumor virus receptor A
VAChT: vesicular acetylcholine transporter
VGAT: vesicular GABA transporter

